# A reliable *in vitro* rumen culture system and workflow for screening anti-methanogenic compounds

**DOI:** 10.1101/2025.03.03.641173

**Authors:** Philip P. Laric, Armina Mortazavi, Ewa Węgrzyn, Kathrin Simon, Pauline S. Rittel, Florian M. Trefz, Benedikt Sabass

## Abstract

Arguably the biggest man-made challenge of the century is to halt the progression of the climate change. Livestock’s methane (CH_4_) emissions, a greenhouse gas with a higher global warming potential than carbon dioxide (CO_2_), represent a prime target for reducing anthropogenic impact. While the reduction of enteric methane emissions through feed additives has been demonstrated, potent and affordable compounds inhibiting methanogenesis in ruminants are hardly established. Reliable methods for reproducible cultivation of the rumen microbiome in the laboratory are an essential tool for the study of methanogenesis.

We have developed a versatile setup that allows for the cultivation of ruminal fluid in a benchtop configuration. This setup combines, miniaturises and improves existing systems. We use everyday labware to create a setup capable of the long term cultivation of primary cultures extracted from the rumen of slaughtered cows. We describe the detailed preparation and cultivation procedure and demonstrate the expected function of the setup. The efficacy of the system is demonstrated through the administration of various concentrations of state-of-the-art methanogenesis inhibitors, including lyophilised *Asparagopsis taxiformis* (AT) powder, bromoform (BF), iodoform (IF), 3-nitrooxypropanol (3-NOP), rapeseed oil and BF dissolved in rapeseed oil. The parameters of the system exhibit a change in accordance with the literature. In comparison with conventional methodologies, our system offers enhanced versatility and ease of use. Furthermore, a novel approach for the quantification of the exhaled gases, CH_4_ and CO_2_, is presented.

## Introduction

The goal of reaching net-zero emissions by the second half of this century was set in the 2015 Paris Agreement (2). Methane (CH_4_) plays a particularly important role for achievement of this goal (3). The atmospheric CH_4_ concentration has tripled since the beginning of the industrial revolution (4). Methane has a relatively short atmospheric half-life of 9.1 ± 0.9 years (5), but a much higher short-term global warming potential (GWP) when compared to carbon dioxide (CO_2_). It is estimated that 1 gram of CH_4_ has the GWP of 34 grams of CO_2_ on a time horizon of 100 years, but 86 grams of CO_2_ over 20 years (6). Consequently the reduction of CH_4_ emissions provides a promising avenue to mitigate global warming.

Livestock production is responsible for approximately 6.38 % of global greenhouse gas (GHG) emissions, equivalent to 3.1 Gt of CO_2_e (CO_2_ equivalents) per year, mostly in the form of CH_4_ and nitrous oxide. This includes enteric fermentation, manure and rice feed (7–9). With global livestock production projected to continue to rise, the reduction of these emissions is of increasing urgency. The digestive system of ruminants contains a complex microbiome consisting of anaerobic bacteria, archaea, protozoa, and fungi. These organisms enable their host to extract nutrients from otherwise mostly indigestible parts of plants. In this symbiosis, the rumen microbiota degrade substrates rich in polymerized structural carbohydrates of otherwise low nutritional value (10). The main fermentation end products are short-chain fatty acids such as propionate, butyrate, acetate, and the gases CO_2_ and hydrogen (H_2_). As increased concentrations of H_2_ inhibit, for example, the redox reaction of nicotinamide adenine dinucleotide (NAD), H_2_ needs to be effectively removed from the system. The most important H_2_ sink is its use as an educt in the metabolism of methanogens in the rumen. Methanogens obtain energy for their growth by the reduction of various substrates, such as CO_2_, to CH_4_ with the aid of H_2_ (11). Suppression of methanogenesis has been shown to result in a redirection of metabolic H_2_ pathways (12,13). For instance *Ruminococcus albus*, one of the major plant cell-wall degrading bacteria, responds to an increase in the partial pressure of H_2_ by a change in the pattern of fermentation products. In the absence of methanogenic archaea, less H_2_ is produced. The normally dominant metabolism of cellulose to butyrate or acetate is then replaced by metabolic pathways that produce less H_2_, such as propionate or valerate (14). As a result of the eructation of CH_4_, ruminants lose approximately 2%-12% of their energy intake. Digestion parameters associated with different levels of CH_4_ production depend on a complex interplay between the microbiome, the animal and the feed. Feed processing techniques used at the farm level, as well as the specific nutrients present in feed formulations, significantly affect CH_4_ production (15). In addition, genetic variation among animals contributes to differences in CH_4_ emissions (16), underscoring the complexity of methanogenesis in ruminants and the necessity of *in vitro* systems for clarifying the mode of action of different feed additives.

There is a wide range of feed additives available to manipulate the rumen microbiome including phytochemicals and essential oils, red algae, nitrates, ionophores, haloalkanes, halogenated sulfonate compounds, nitrooxy compounds, bacteriocins, addition of acetogens, and methods to control the protozoan population (17). To provide context for this work, we briefly review a few prominent anti-methanogenic additives.

One extensively studied group of inhibitors of microbial methanogenesis are polyhalogenated compounds that can be characterized as CH_4_ analogs. Initial studies investigated the use of carbon tetrachloride and chloroform (18). Means to reduce methanogenesis were proposed in order to protect sheep from the harmful effects of *Heliotropium europaeum*, a plant that contains hepatotoxic pyrrolizidine alkaloids, whose degradation is favored by elevated H_2_ levels (19–21). Further polyhalogenated, anti-methanogenic compounds include chloral hydrate (22), tribromoacetaldehyde, bromoform (BF), and carbon tetrabromide (23–26). A recent study with iodoform (IF) showed good anti-methanogenic activity, but as a side effect, milk yield and feed intake were reduced. Furthermore, alterations of several metabolic markers were indicative of the presence of a negative energy balance. Reduced feed intake may result directly from the administered IF, an excess of iodine, as iodine was also supplemented, or a refusal of the diet because of the presumed strong taste of IF during rumination (27).

Promising natural anti-methanogenic feed additives are red algae, in particular *Asparagopsis taxiformis* (AT). The the main active agent in AT is bromoform. Supplementation of AT can result in up to 80% reduction of CH_4_ production and a reduction in feed intake with no apparent negative effect on average daily weight gain (28,29). However, high concentrations of BF are considered toxic and can be harmful if ingested. There are concerns about negative effects on the environment and the ozone layer, as excessive production and feeding of AT could lead to increased emissions of halocarbons (30). Furthermore, supplementation of BF containing compounds can lead to the formation of volatile dibromomethane and bromomethane in the rumen (31).

An entirely different group of anti-methanogenic compounds are nitro esters. The well-studied small molecule 3-Nitrooxypropanol (3-NOP) inhibits Methyl-Coenzyme-M-Reductase (MCR). This enzyme catalyzes the final step in the formation of CH_4_ by combining the H_2_ donor coenzyme B and the methyl donor coenzyme M, see Fig 1 (32). Suppression of methanogenesis is thought to occur through specific binding of 3-NOP to the active center of MCR, where it oxidizes Ni(I) (33). Other recently discovered anti-methanogenic compounds containing nitro esters include N-[2-(nitrooxy)ethyl]-3-pyridinecarboxamide (NPD), 2,2-dimethyl-3-(nitrooxy) propanoic acid (DNP) and nitroglycerin. Notably, all of these substances show distinct inhibitory properties due to their molecular structures, with nitroglycerin, which contains three nitrooxy moieties, exhibiting the strongest inhibitory activity (34).

**Figure 1:**
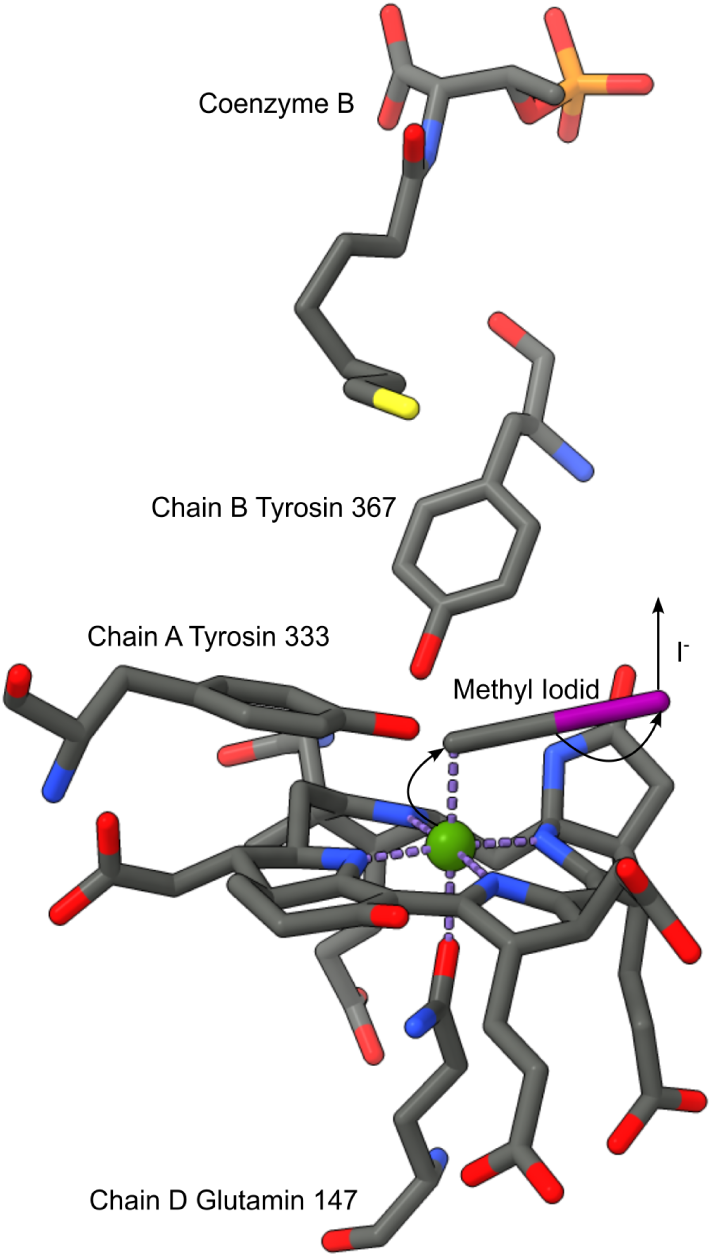
X-ray-cristallography model of the active center of the MCR; cofactor F430 with methyl iodide and Coenzyme B. Grey: carbon; blue: nitrogen; red: oxygen; yellow: sulfur; pink: iodide; green: nickel(III) (1).

There is a considerable amount of published research on ruminal fermentation, which typically focuses on evaluating feed value and digestion efficiency. *In vitro* experiments are frequently done with cultures from rumen inoculi, see Fig. 2. Rumen microbiome cultivation methods can be categorized into two different approaches, namely, short-term batch cultivation and long-term rumen simulation techniques (RUSITEC), both of which will be discussed below. The batch culture system is characterized by its ease of use with possible high probe throughput. Batch culture systems have been designed to evaluate the short-term effects of a compound on fermentation parameters (18,20). The integration of an automated gas accumulation recording and release system (ANKOM) has further expanded the utility of laboratory flasks in batch experiments (28). For operation, ruminal fluid is diluted with buffer and transferred to vessels prepared with different compounds of interest and the substrate. The vessels are subsequently incubated at 39°C. Typically, the duration of these batch experiments is up to three days, until the culture has fully depleted the nutrients from the substrate. Recently, automated high-throughput systems in well plate format have become available (35,36), with the potential to miniaturize the batch setup to a 96-well plate format, thereby enhancing the throughput. However, miniaturization imposes constraints on the parameters that can be analyzed, confining them to gas production parameters and the nutrients supplied. The fundamental limitation of the batch culture system is its incapacity to sustain a stable microbiome, thus preventing long-term experiments to assess the effects on the microbiome and the consistency of the treatment effects. The decline in certain species observed during the experiments is indicative of a diminished quality of the measurements (37). RUSITEC setups are designed to simulate the conditions of the rumen in more detail and consist of a series of multiple vessels that mimic the rumen environment. These setups maintain a more stable microbiome that can closely resemble the original rumen inoculum (38). Despite the enhanced stability of the microbiome in rumen simulation fermenters, a significant loss of protozoal biomass has been documented (39). In recent years, various systems capable of sustaining the whole microbiome during prolonged cultivation have been designed. Examples of such state-of-the-art systems include semi-continuous single-flow fermenters(40), continuous dual-flow fermenters that retain some of the solids effluent by filtration (41) and intermittent single-flow systems that stratify the fermenter contents by slow or intermittent agitation (42). The semi-continuous single-flow fermenter represents the most prominent approach. In this configuration, the diet is stored in a permeable bag within the system and a system of tubes and pumps facilitates the constant exchange of culture and fresh buffer. Regular feeding is achieved by opening the system and the replacement of the bag inside (40). From a practical point of view, a limitation of the RUSITEC approach is the low probe throughput, because the complicated and expensive systems cannot be easily multiplexed.

**Figure 2:**
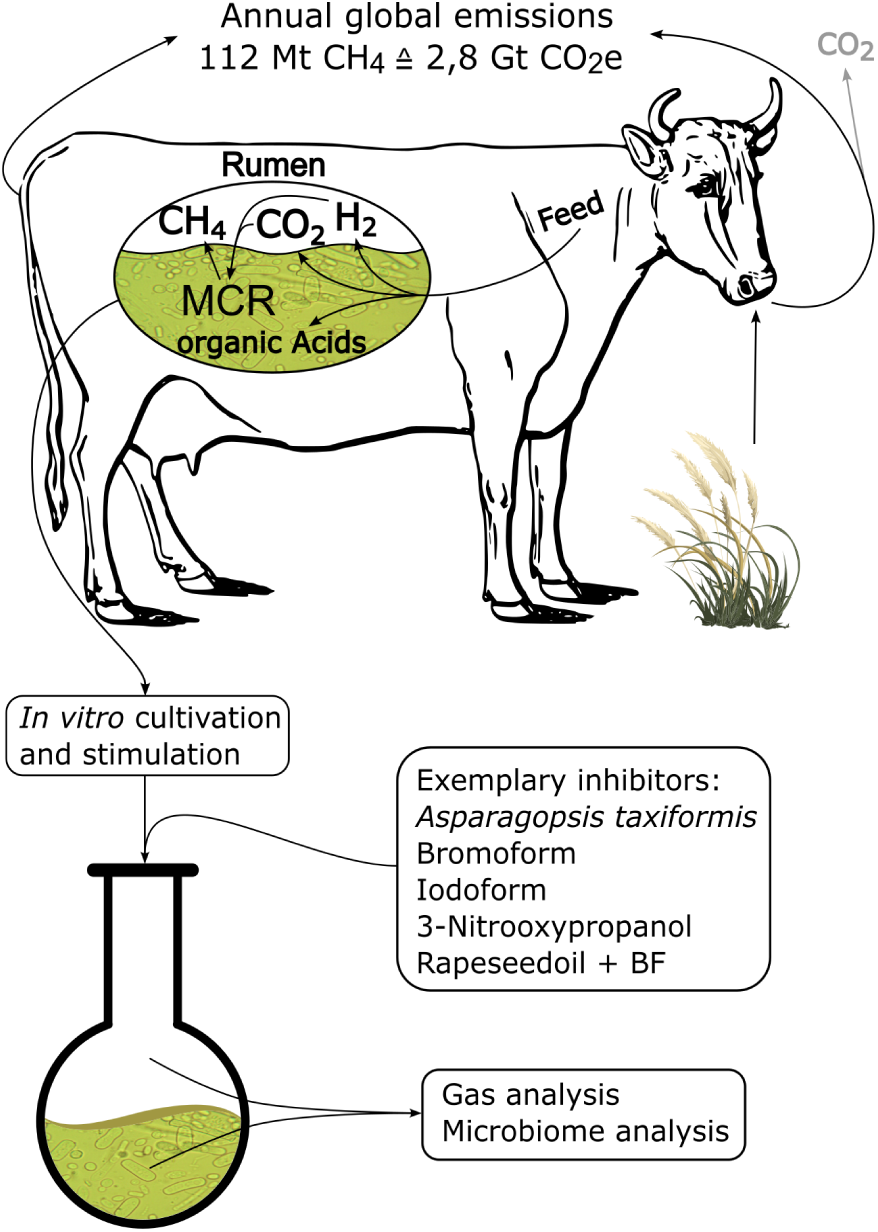
Methanogenesis in the cow rumen and its yearly methane exhaust. Overview of workflow for inhibitor development *in vitro*.

We report here the design of a reliable and easy-to-use, semi-continuous rumen simulation system that is optimized for testing anti-methanogenic supplements. Detailed assembly instructions are provided and the setup is designed to be built at minimal cost. To achieve this, we used everyday laboratory equipment that was crafted into an anaerobic incubation apparatus. Batch cultivation approaches were combined with RUSITEC approaches to create a continuous batch system. The integrated inlets and outlets facilitate daily probing and feeding without disturbing the system, as well as stimulations with compounds of interest. To demonstrate the use of the setup, we compared the CH_4_, CO_2_ and residual gas productions in the presence of different concentrations of various anti-methanogenic supplements. In detail, we compared AT (28), BF (43–45), IF (23,27), 3-NOP (46–49), rapeseed oil and BF dissolved in rapeseed oil (50,51), while tracking shifts in the microbiome. Thereby, we provide a so-far lacking direct comparison between the most studied compounds (52,53) and additive formulations (54).

## Material&Methods

### Biological materials

#### *In vitro* cultivation

- Rumen fluid and solids extracted from four different rumen

### Reagents

#### Gas measurements

- CH_4_ 2,5% in N_2_ (ALL-IN-GAS, Munich, Germany)
- CH_4_ 30% in N_2_ (ALL-IN-GAS, Munich, Germany)
- CO_2_ (2.5, Linde, Pullach, Germany)
- N_2_ (5.0, Linde, Pullach, Germany)

#### *In vitro* cultivation

- (NH_4_)_2_SO_4_ (≥99,5%, Carl Roth, Karlsruhe, Germany)
- Acetic acid (≥99%, Carl Roth, Karlsruhe, Germany)
- Butyric acid (p.s., Sigma-Aldrich, Steinheim, Germany)
- CaCl_2_ (≥98%, Carl Roth, Karlsruhe, Germany)
- Cellobiose (≥98%, Carl Roth, Karlsruhe, Germany)
- CO_2_ (2.5, Linde, Pullach, Germany)
- DL-2-Methyl butyric acid (≥98%, Thermo Fisher Scientific, Waltham, Massachusetts, USA)
- Glucose (≥98%, Carl Roth, Karlsruhe, Germany)
- Glycerol (≥99,5%, Sigma-Aldrich, Steinheim, Germany)
- H_2_O desalted (dH_2_O) (GENO-OSMO-X, Grünbeck, Höchstädt, Germany)
- HCl (37%, Carl Roth, Karlsruhe, Germany)
- Hemin chlorid (≥98%, Carl Roth, Karlsruhe, Germany)
- High-fiber silage (Ströh, Hamburg, Germany)
- iso-Butyric acid (≥99%, Carl Roth, Karlsruhe, Germany)
- iso-Valeric acid (p.s., Sigma-Aldrich, Steinheim, Germany)
- K_2_HPO_4_ (≥98%, Sigma-Aldrich, Steinheim, Germany)
- KCl (≥99%, Carl Roth, Karlsruhe, Germany)
- KH_2_PO_4_ (p.a., Carl Roth, Karlsruhe, Germany)
- KOH (≥85%, Carl Roth, Karlsruhe, Germany)
- L-cysteine-HCl * H_2_O (≥98,5%, Carl Roth, Karlsruhe, Germany)
- Maltose (≥97%, Carl Roth, Karlsruhe, Germany)
- MgCl_2_ (≥99%, Carl Roth, Karlsruhe, Germany)
- MgSO_4_ (≥99%, Carl Roth, Karlsruhe, Germany)
- n-Valeric acid (≥99%, Thermo Fisher Scientific, Waltham, Massachusetts, USA)
- Na-resazurin (Alfa Aesar, Haverhill, Massachusetts, USA)
- Na_2_CO_3_ (VWR, Darmstadt, Germany)
- Na_2_HPO_4_ (≥99%, Carl Roth, Karlsruhe, Germany)
- Na_2_S * 9 H_2_O (≥98%, Thermo Fisher Scientific, Waltham, Massachusetts, USA)
- NaCl (≥99,5%, Carl Roth, Karlsruhe, Germany)
- NaHCO_3_ (≥99,5%, Carl Roth, Karlsruhe, Germany)
- NaOH (≥98%, Carl Roth, Karlsruhe, Germany)
- NaOH (p.a., Carl Roth, Karlsruhe, Germany)
- Propionic acid (≥99%, VWR, Darmstadt, Germany)
- Starch, soluble (p.a., Carl Roth, Karlsruhe, Germany)
- Yeast extract (for cellculture, Carl Roth, Karlsruhe, Germany)

#### Microbiome analysis

- Agarose-LE (Biozym Scientific, Oldendorf, Germany)
- AMPure XP
- CH_3_COOH (≥99%, Carl Roth, Karlsruhe, Germany)
- Deoxynucleotide solution (New England Biolabs, Frankfurt am Main, Germany)
- DNA extraction kit (ExtractME, BLIRT S.A., Gdańsk, Poland)
- DNA ladder 1 kb (New England Biolabs, Frankfurt am Main, Germany)
- EDTA (Merck, Darmstadt, Germany)
- Ethanol (≥99,5%, Carl Roth, Karlsruhe, Germany)
- Gel leading dye blue 6x (New England Biolabs, Frankfurt am Main, Germany)
- GelRed® Nucleic acid stain 10000x (New England Biolabs, Frankfurt am Main, Germany)
- Isopropanol (≥99,8%, Sigma-Aldrich, Steinheim, Germany)
- NaCl (≥99,5%, Carl Roth, Karlsruhe, Germany)
- NH_4_CH_3_COO (≥99%, Carl Roth, Karlsruhe, Germany)
- Q5® High-fidelity DNA polymerase (New England Biolabs, Frankfurt am Main, Germany)
- SDS (≥99%, Carl Roth, Karlsruhe, Germany)
- Tris-HCl (≥99%, Sigma-Aldrich, Steinheim, Germany)
- Tris-OH (≥99%, Sigma-Aldrich, Steinheim, Germany)

#### Stimulants

- 3-Nitrooxypropanol (≥99%, synthesised and verified by Ewa Wegrzyn, Chemical and Pharmazeutical Faculty, LMU-Munich, Butenandtstraße 5, 81377 Munich)
- Bromoform (≥99%, Merck, Darmstadt, Germany)
- Ethanol (≥99,8%, Carl Roth, Karlsruhe, Germany)
- Freeze-dried AT powder ( 4.5 g BF/g powder, Volta-Greentech, Solna, Sweden)
- Iodoform (99%, Thermo Fischer, Massachusetts, USA)
- Rapeseed oil (native, Alnatura GmbH, Darmstadt, Germany)

### Equipment

#### General equipment

- 8-Channel pipette (Ergonomic High Performance multichannel pipette, VWR, Darmstadt, Germany)
- Autoclave (HG-133, HMC-Europe, Tüßling, Germany)
- Epoxy resin (Epoxyplast 100 P, DIPON.DE, Dortmund, Germany)
- Falcon tubes (Sarstedt, Nümbrecht, Germany)
- Glue gun (TC-GG 30, Einhell Germany, Landau/Isar, Germany)
- Luer-lock adaptor female/female (neoLab, Heidelberg, Germany)
- Luer-Lock barbed adaptor 6mm on female and male (neoLab, Heidelberg, Germany)
- Luer-Lock valve male/female (Discofix®, B.Braun, Melsungen, Germany)
- Magnetic stirrer (ROTILABO®MH-15, Carl Roth, Karlsruhe, Germany)
- Pipettes 2 - 1000 µL (Eppendorf, Hamburg, Germany)
- Reaction tubes (Eppendorf, Hamburg, Germany)
- Serological pipettes (Sarstedt, Nümbrecht, Germany)
- Silicon tubes Ø 6mm (ESSKA, Hamburg, Germany)
- Vaseline (Heinrich Hagner GmbH, Freudenstadt, Germany)

#### Gas measurements

- 3 way connector 7 mm (OBI, Wermelskirchen, Germany)
- Fourier-transform infrared (FT-IR) spektrophotometer (SpektrumTwo, Perkin Elmer, Massachusetts, USA)
- Gas cuvette (Storm10 cm pyrex, Specac Ltd., UK)
- Needles 0,80 x 120 mm (Sterican®, B.Braun, Melsungen, Germany)
- Reservoir bottle with cap 150 mL (VWR, Darmstadt, Germany)
- Rubber stoppers 5×9×20 (Carl Roth, Karlsruhe, Germany)
- Syringes 200 and 300 mL (Romed, Wilnis, Netherlands)
- Windows for the gas cuvette (CaF_2_ Window-Pair, Specac Ltd., UK)

#### *In vitro* cultivation

- 1 L Erlenmeyer (DWK Life Sciences, Mainz, Germany)
- 10 L Polypropylene bucket (VWR, Darmstadt, Germany)
- Autoclave ()
- Balloons (Latex, various, Amazon)
- Blender 2000W (Homegeek, Huanan, China)
- Boning knife (F. Dick, Deizisau, Germany)
- Bottletop filters Ø 0.22 µm (VWR, Darmstadt, Germany)
- Cheesecloth (Päckärdi, Huddersfield, UK)
- Cool packs (Various)
- Fabric hose 8 mm (ESSKA, Hamburg, Germany)
- Immersion blender (Philips 5000 series, Philips, Amsterdam, Netherlands)
- Incubation hood (Medium, neoLab, Heidelberg, Germany)
- Ladle (IKEA, Delft, Niederlande)
- Orbital shaker (WS-1500, Wiggens, Straubenhardt, Germany)
- pH Meter (accumet AE150, Fischer Scientific, Schwerte, Germany)
- Polypropylene bucket 1 L (Berry Global, Evansville, USA)
- Sieve 1 mm mesh (IKEA, Delft, Niederlande)
- Syringe filter 0.22 µm (TPP, Trasadingen, Switzerland)
- Syringes 100 mL (Romed, Wilnis, Netherlands)
- Vaseline (Heinrich Hagner GmbH, Freudenstadt, Germany)
- Waterbath (WB-12, Phoenix Instrument, Garbsen, Germany)

#### Microbiome analysis

- Bead beater (Tissue Lyser MM300, QUIAGEN, Hilden, Germany)
- Centrifuge (Fresco™ 21 Microcentrifuge, Thermo Fisher Scientific, Waltham, Massachusetts, USA)
- Gel Caster (Bio-Rad Laboratories, Hercules, California, USA)
- Gel comb (10-Well comb, Bio-Rad Laboratories, Hercules, California, USA)
- Gel documentation system (Vilber Lourmat, Biovision 3000 WL, Eberhardzell, Germany)
- Gel tray (7×8 cm UV-transparent mini-gel tray, Bio-Rad Laboratories, Hercules, California, USA)
- Glass beads 0.1 mm (Carl Roth, Karlsruhe, Germany)
- Horizontal electrophoresis system (Mini-Sub Cell GT Cell, Bio-Rad Laboratories, Hercules, California, USA)
- Imaging chamber (Darkroom – CN3000, PEQLAB, Erlangen, Germany)
- Neodymium magnets (EarthMag, Dortmund, Germany)
- PCR tube strips (Eppendorf, Hamburg, Germany)
- Power supply (PowerPac™ Basic, Bio-Rad Laboratories, Hercules, California, USA)
- Rack for PCR tubes (neoLab, Heidelberg, Germany)
- Thermocycler (Vapo Protect Mastercycler® Pro, Eppendorf, Hamburg, Germany)
- ThermoMixer® C (Eppendorf, Hamburg, Germany)
- UV-Vis cuvettes (PMMA, VWR, Darmstadt, Germany)
- UV-Vis spectrophotometer (Nanodrop1000, Thermo Fisher Scientific, Waltham, Massachusetts, USA)
- Vortexer (Vortex-Genie™ 2, Scientific Industries, Bohemia, New York, USA)
- Zirconia beads 0.5 mm (Scientific Industries, Bohemia, New York, USA)

#### Setup design

- 3 D Printer (PRUSA i3 MK3, Prusa Research, Prag, Czech Republic)
- 500 mL GL45 laboratory bottles (DURAN®pressureplus, DWK Life Sciences, Mainz, Germany)
- 500 mL Spout bags (Daklapack, Lelystad, Netherlands)
- Bromobutyl plug seal closure (DURAN®, DWK Life Sciences, Mainz, Germany)
- Duct tape (3M, Neuss, Germany)
- Engraving tool (Dremel 3000, Dremel Europe, Breda, Netherlands)
- High-speed milling set (Dremel Europe, Breda, Netherlands)
- Open topped screw cap GL45 (DURAN®, DWK Life Sciences, Mainz, Germany)
- Polyethylenterephthalat-Filament (Polymaker, Houten, Netherlands)
- Toolbox (Conmetall Meister, Celle, Germany) optional:
- Exsiccator (554, Kartell, Novoglio, Italy)
- Incubator (INCU-Line® IL56, VWR, Darmstadt, Germany)
- Resin colour or pigment (DIPON.DE, Dortmund, Germany)
- Vacuum pump (MZ 2 NT, Vacuubrand, Wertheim, Germany)
- Waterbath (WB-12, Phoenix Instrument, Garbsen, Germany)

### Software

#### Gas measurements

- Spectrum (Perkin Elmer, Massachusetts, USA)
- Spectrum Quant (Perkin Elmer, Massachusetts, USA)

#### Microbiome analysis

- DADA2 package (v.1.14) (55)
- DNA-Quantification (Nanodrop1000, Thermo Fisher Scientific, Waltham, Massachusetts, USA)
- Gel visulalisation (VISION-CAPT, Vilber Lourmat, Eberhardzell, Germany)
- R-Studio 2022.02.2+485 (PBC, Boston, USA)

#### Setup design

- Prusa Slicer (Prusa Research, Prag, Czech Republic)
- Solidworks 2017 (Dassault Systèmes, Stuttgart, Germany)

#### Evaluation

- UCSF ChimeraX v. 1.9 (56)
- GraphPadPRISM 5 (GraphPadSoftware, Boston, Massachusetts, USA)
- Inkscape 1.4 (Inkscape Community, Brooklyn, New York, USA)
- LibreOffice 24.8.4 (The Document Foundation, Berlin, Gemany)

### External services

#### Microbiome analysis

- Sequencing service (GENEWIZ Germany GmbH, Leipzig)

### Reagent setup

#### Setup design

Epoxy resin: mix two parts of component A with one part of component B. Stir thoroughly. Note: Use alcohol ink or pigment to ensure proper mixing of the components. To shorten the pot life of the resin and enhance mixability, warm the components in a water bath. For the removal of bubbles the container with the mixed resin can be evacuated, removed with a lighter flame or with isopropanol. It is good practice to roughen up the surfaces, which are in contact with the resin to ensure a strong bond. Make sure to wait long enough for the resin to cure.

#### *In vitro* cultivation

Cellobiose stock: Dissolve 5 g of cellobiose in 100 mL dH_2_O and sterilise by filtration. Store at room temperature for several months.

Cysteine-HCl stock: Dissolve 1 g of L-cysteine-HCl * H_2_O in 50 mL of dH_2_O and sterilise by filtration. Store at room temperature for several months.

Glucose stock: Dissolve 20 g of glucose in 100 mL of dH_2_O and sterilise by filtration. Store at room temperature for several months.

Hemin solution: Dissolve 50 mg of Hemin chlorid in 1 mL 1 M NaOH and fill up to 100 mL with dH_2_O. Store at 4-8 °C for several months.

Maltose stock: Dissolve 20 g of maltose in 100 mL of dH_2_O and sterilise by filtration. Store at room temperature for several months.

Mineral Solution: Dissolve 3 g of KH_2_PO_4_, 6 g of NaCl, 3 g of (NH_4_)_2_SO_4_, 0.6 g of CaCl_2_ and 0.61 g of MgSO_4_ in 1 L of dH_2_O and sterilise by filtration. Store at room temperature for several months.

Note: For stability, add one to two drops of HCl.

Na-resazurin solution: Dissolve 10 mg of Na-resazurin in 10 mL dH_2_O and sterilise by filtration.

Na_2_S solution: Dissolve 8.5 g of Na_2_S * 9 H_2_O in 85 mL of dH_2_O and dispense into 8.5 mL alliquotes. Store them in a tightly closed container at -20°C.

Rumen bacteria medium: Prepare the rumen bacteria medium by dissolving 0.3 g of K_2_HPO_4_, 0.5 g of yeast extract, 0.5 g of Glycerol, 0.5 g of starch soluble, 38 mL of mineral solution, 1.6 mL of Na-resazurin solution, 3.1 mL of volatile fatty acids solution and 2 mL of hemin solution in 926.4 mL of dH_2_O. Autoclave prior to completion with 2.5 mL of glucose stock, 2.5 mL of maltose stock, 10 mL of cellobiose stock, 12.5 mL of L-cysteine stock, 4 g of Na_2_CO_3_ and 2.5 mL of Na_2_S stock. If necessary, adjust the pH of the complete media with KOH or by sparging with CO_2_ to 6.7-6.8 (57).

Saliva buffer: Dissolve 22.2 mg of CaCl_2_, 28.58 mg of MgCl_2_, 9.8 g of NaHCO_3_, 4.63 g of Na_2_HPO_4_, 0.47 g of NaCl and 0.57 g of KCl. If necessary adjust the pH with NaOH or by sparging with CO_2_ to 8.2 (58).

Volatile fatty acids solution: Mix together 548.5 mL of acetic acid, 193.5 mL of propionic acid, 129 mL of butyric acid, 32.25 mL of isobutyric acid, 32.25 mL of DL-2-methylbutyric acid, 32.25 mL of valeric acid and 32.25 mL of isovaleric acid. Store at room temperature.

### Procedure

#### Gas measurements

FT-IR assembly

1. Set up the spectrophotometer and the gas cuvette according to the manual.
2. Mill a sealing ring for the 3-way connector out of a rubber stopper and put both into the inlet position.
3. Repeat for the female luer-lock adaptor and a rubber stopper for the outlet position.
4. Now attach one luer-lock valve to each opening and check if the system is airtight.
5. Attach another luer-lock valve to a short piece of silicon tube via a male luer-lock adaptor.
6. One luer-lock valve on the 3-way connector is for the N_2_ supply, the other is the injection port and the N_2_ tap.
7. Take the reservoir bottle and drill two holes into the screwcap.
8. Penetrate a rubber stopper with several needles, cut off the luer adaptor and plug it into the silicon tubing. Note: Make sure the cut does not obstruct the canule.
9. Fix the tube with the needles to one hole of the screwcap. The needles should reach to the bottom of the flask.
10. Attach another tube to the second hole. 3D-Print a cylindrical mold and glue it to the screwcap. Pour epoxy into the mold and let it cure.
11. Fill tap water into the bottle until all needles are submerged.
12. Lead the exhaust tube to a ventilation pipe. Syringe assembly
13. Take a 200 mL and a 300 mL Syringe and cut off the tip, that the female end of a luer-lock valve fits into and does not obstruct the piston of the syringe.
14. 3D-Print two cylindric sections that leave room for the valve movement and have sufficient attachment area.
15. Seal the female ends of the luer-lock valves with vaseline and apply vaseline to the syringe piston aswell.
16. Insert the luer-lock valve into the syringe tip and move the piston into exhaust position.
17. Hot-glue the mold into position, cast the epoxy and let it cure. Note: Move the valve into closed position to prevent obstruction of the valve by the epoxy.
18. Check for leakage. Generation of the calibration curve for N_2_ and CO_2_
19. Start the FT-IR spectrophotometer and the spectrum software. Insert the assembled gas cuvette into the lightpath and attach the N_2_ gas line to the gas inlet of the cuvette.
20. Flush the gas cuvette and record the background
21. Measure at least three spectra of one concentration. Note: To prepare diluted gas standards, draw calibration gas into the 300-mL syringe. Adjust the volume based on the dilution factor. Exhaust the gas through a needle into a water-filled vessel and wait for the pressure to equilibrate to precisely measure the volume. Then take the 200-mL syringe and draw the needed amount of N_2_ gas to fill the 300-mL syringe to 300 mL. Consequently, transfer the gas into the 300-mL syringe with a female/female adaptor.
22. Save the spectra and load them into the spectrum quant software to generate the quantification method according to the manual.
23. Load the generated methods into the spectrum software. Evaluation of the *in vitro* gas production
24. After the feeding, close both luer-lock valves and exchange the gas bag. Note: The gas bags can now be stored until evaluation either at room temperature or at -20°C.
25. Shortly freeze the filled gas bags to reduce the gas humidity.
26. Start the spectrum software, flush the gas cuvette with N_2_ and record the background.
27. Draw 300 mL of gas from the gas bags into the 300-mL syringe. Note: If the gas production was less than 300 mL, go to the next 50 mL step and substitute the missing volume with N_2_.
28. Measure the residual gas content of the gas bag with the 200-mL syringe and note the total gas production.
29. Inject 300 mL of gas into the gas cuvette via the injection port.
30. Wait for the pressure to equilibrate and record the measurement.
31. After all samples are measured, run the evaluation methods to quantify the gas contents.
32. Proceed with data evaluation

#### *In vitro* cultivation

Rumen culture studies were conducted for 11 days. Stimulation with additives was performed on the fourth day. A long-term experiment was conducted where rumen fluid was cultured for 23 days with a second stimulation on day 16.

Preparations

1. Dry the silage and blend it for 2 minutes. Then sieve it through a 1 mm mesh and return the residues into the blender.
2. Prepare two feeding syringes.
3. Cut off the tip of the syringes but 1 cm.
4. Take a 50 mL serological pipette and cut the tip to the size of your silicon tubes.
5. Deburr the edges with a cutter knife.
6. In addition, trim the other end to expose the full width of the pipette.
7. 3D-print a cylindrical mold and fix it with hot glue to the syringe.
8. Balance the serological pipette on the syringe and cast epoxy into the mold.
9. Check for leakage. Sample collection
10. Fill the 1 L bucket with tap water and prewarm it with the cool packs to 55°C the day prior to the slaughterhouse visit.
11. In the morning of the slaughterhouse visit, complete the rumen media and put it into the waterbath at 39°C, weigh 2 g of ground silage and add 10 ceramic loops per culture bottle and put the bottles into the incubator at 39°C.
12. At the slaughterhouse collect rumen fluid and content right after evisceration, through an incision of the rumen of four randomly selected cattle.
13. Perform the subsequent steps of homogenisation, straining and the preparation for transport as previously described (28).
14. Prepare the inoculum by adding one part of rumen fluid to four parts of rumen media.
15. Pour the culture into the prewarmed culture bottles and close them with the assembled screwcaps.
16. Purge the culture with pure N_2_ for 2 minutes.
17. Put the bottles into the orbital shaker under the incubation hood and attach the gas bags.
18. Set the orbital shaker to an intermittent shaking program of five minutes shaking followed by 5 minutes of pause and set the incubation hood to 39°C. Daily feeding
19. About half an hour prior to the feeding, weigh in 2 g of ground silage per culture, add 50 mL of saliva buffer (58) per culture and gently bubble it with nitrogen under vigorous stirring.
20. Take the bottles out of the shaker and incubation hood one by one.
21. Remove the plug, swirl once and immediately withdraw 50 mL of culture and either take samples or throw them away.
22. With the other pipette draw 50 mL of the feeding mix and inject them into the flask.
23. Insert the plug and exchange the gasbag and put it back into the incubator Stimulation
24. Prepare the stimulants the day before stimulation. Note: It is best to dissolve the stimulant, as it facilitates the handling. For the addition of solids, remove the bottom of an 1.5 mL eppi and cover it with parafilm, then weigh in the stimulant. Now the eppi can be attached to the silicon tube and the feeding solution can be flushed through the eppi.
25. Stimulate the cultures on the fourth day of cultivation with 0.5 mL to 1.5 mL of liquid stimulant stock solution or any amount of solid stimulant.
26. Add the compounds prior to the injection of the feeding mix to wash all of the stimulant in.

#### Microbiome analysis

Preparations

1. Prepare the magnetic stand for the AMPure XP beads. Use hot glue to fix one magnet each in north-south direction between two holes under the PCR rack. Sample collection
2. From the daily feed waste, take 2 x 2 mL of samples with a cut-off pipette tip to also transfer solid contents.
3. the solids from the fluid by a centrifugation step at 10 000 x g for 5 min at 4°C.
4. the supernatant and freeze the pellett at -80°C until further processing. Repeated bead-beating + column (59)
5. Thaw the pelletts on ice and add 0.3 g of 0.1 mm beads and 0.1 g of 0.5 mm beads.
6. Homogenise the pelletts with the bead beater at maximum speed for 3 min.
7. Incubate at 70°C for 15 min with gentle shaking every 5 min in the thermomixer.
8. Centrifuge at 16000 x g at 4°C for 5 min.
9. Transfer the supernatant into a new eppi. Optional: Add another 300 µL lysis buffer to the pellett and repeat steps 5-8 if needed.
10. Add 200 µL of 10 M NH_4_CH_3_COO (260 µL if the optional step was carried out), mix well and incubate on ice for 5 min.
11. Centrifuge at 16000 x g at 4°C for 5 min.
12. Transfer 1 mL of the supernatant to a new 2 mL eppi, add the same volume of ice cold isopropanol, mix well and incubate on ice for 30 min.
13. Centrifuge at 16000 x g at 4°C for 5 min.
14. Discard the supernatant and wash the nucleic acid pellett with ice cold 70% ethanol and dry the pellett.
15. Dissolve the nucleic acid pellett in 150 µL of elution buffer from the kit.
16. Proceed with the DNA cleasnup as described in the manual of the kit.
17. Store the DNA solutions at -20°C until further processing. Polymerase chain reaction (60)
18. Prepare a PCR mix according to the manual of the manufacturer.
19. For the amplification of the 16S small subunit ribosomal RNA fragments following primers were used:
 

**Table 1:**
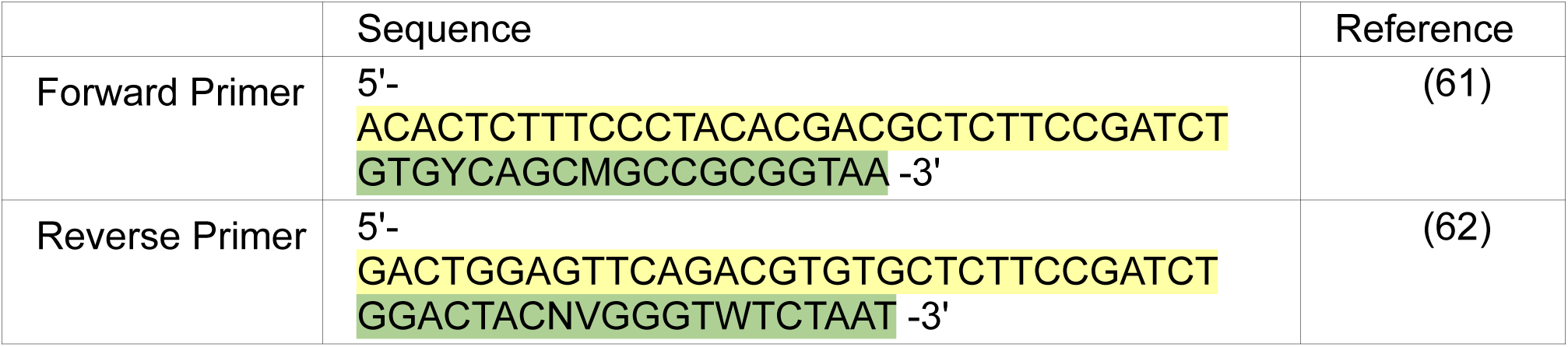
List of primers used for amplification of the 16S small subunit ribosomal RNA fragment subsequent sequencing. The primers are annotated in 3’-5’ direction. In yellow, the sequencing adaptor; in green, the degenerated binding sequence.
20. Seal the individual reaction tubes and perform the PCR with the following settings:
  - 95°C for 3 min
  - 35 cycles of:
    - 95°C for 30 s
    - 55°C for 30 s
    - 72°C for 30 s
  - 72°C for 5 min
  - Hold at 4°C PCR purification
21. Follow the steps of the AMPure XP manual and use the prepared magnetic stand.
22. Check the concentration on the Nanodrop1000 and dilute the concentration to 20 ng/µL
23. Send the tubes to the external sequencing service and wait for the data. Data evaluation (63)
24. Trim the raw sequencing data, then process it in R-Studios according to the exemplary online workflow. In summary:
  - remove the primers
  - quality filter, error correct and merge the reads
  - remove chimeras
  - assign the reads with the DADA2 package with following parameters
    - left and right side trimming at base 20
    - right-side read truncation at base 210 for forward reads
    - left-side read truncation at base 210 for reverse reads
    - default settings otherwise
  - use the SILVA v138 Database to assign the taxonomy
  - generate graphs and tables

#### Setup design

The main vessel consists of a 500 mL GL45 pressure plus laboratory bottle. Screwcap assembly, see Fig. 3.

1. To construct the plug, drill two holes, approximately 5 mm in diameter, into two opposing cavities of the bromobutyl stopper.

**Figure 3:**
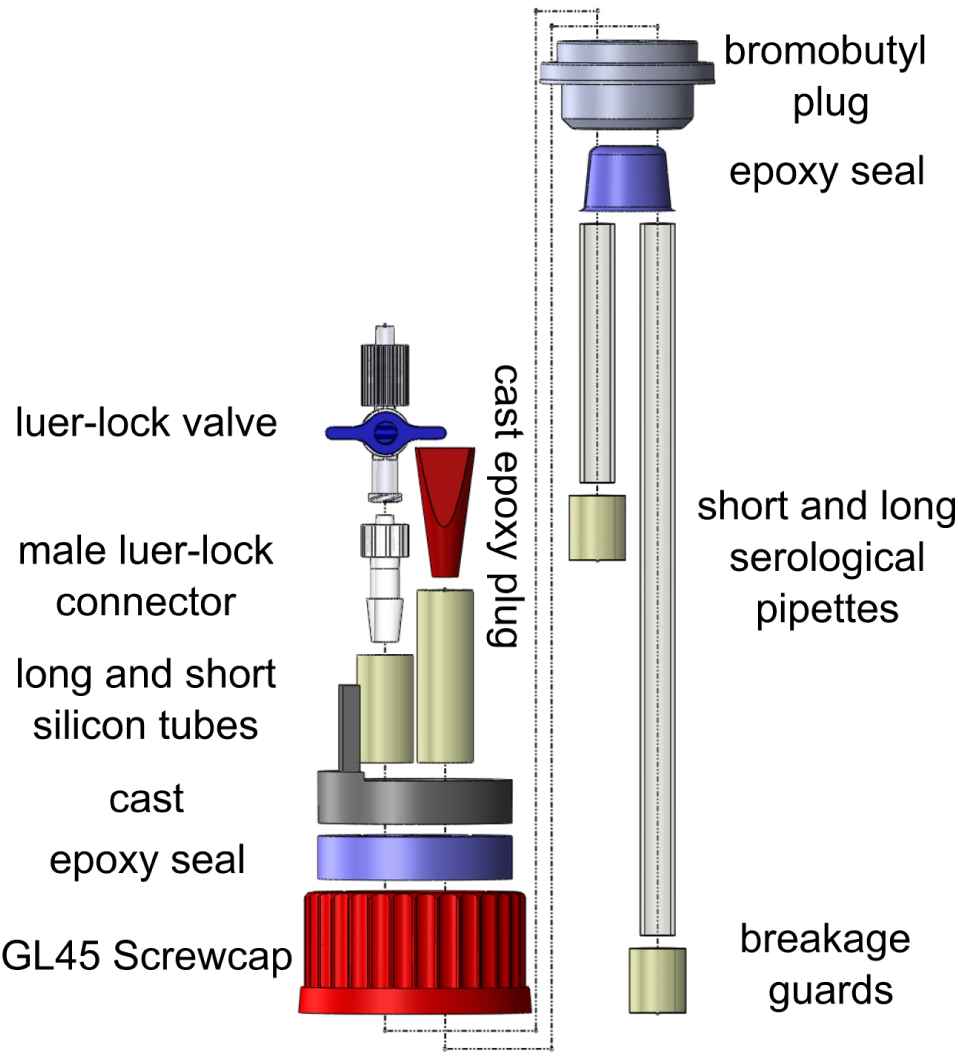
Exploded view of the screwcap assembly. CAUTION The drilling is mostly done by friction, therefore hot flakes of the rubber will fly around
2. Cut two 5 mL serological pipettes with the circular saw for the engraving tool to a length of 165 mm and 60 mm to yield a long and a short piece, respectively. Deburr the edges with a cutter knife.
3. Cut four pieces of silicon tubing to a length of 1, 1, 2.5, and 4 cm.
4. Insert the long pipette piece through the first hole so that it reaches close to the bottom of at the top. Put a 1 cm long piece of silicon tube on the lower end as a breakage guard and a 2.5 cm long piece on the upper end.
5. The short piece should just penetrate the plug for about 2 cm and protrude 1 cm on top aswell. Put on a 1 cm long piece of silicon tubing over the lower ends as breakage guard. The upper end gets attached to the 4 cm long silicon tube.
6. Ensure that the lower part of the bromobutyl stopper is not warped and that the pipettes stand straight, then cast it with epoxy.
7. Draw a cast for on top of the screwcap with the diameter of the screwcap and a height of 1 cm. The reinforcement for the long silicon tube is optional.
8. Fix the mold with hotglue and seal all gaps. Cast the mold with epoxy and let it cure.
9. Stick the male luer-lock adaptor into the long silicon tube and secure the luer-lock valve on it
10. For the short silicon tube, either buy a fitting hard rubber stopper or draw a mold for it and cast it with epoxy a7nd plug it in.
11. Screw the finished cap onto the glass bottle and check for leakage. Note: Apply a little pressure to the bottle and submerge in water. Gasbag assembly, see Fig. 4.
12. Take a spoutbag and reinforce all edges and kinks with duct tape.
13. Cut the spout off.
14. 3D print a female luer-lock adaptor that fits in the hole of the spoutbag.
15. Attach a luer-lock valve and hotglue the adaptor into place and seal it off.
16. Also print a cast for around the spout and the Luer-lock valve that allows for manipulation of the valve.
17. Pour the mold with epoxy
18. Check for leakage

**Figure 4:**
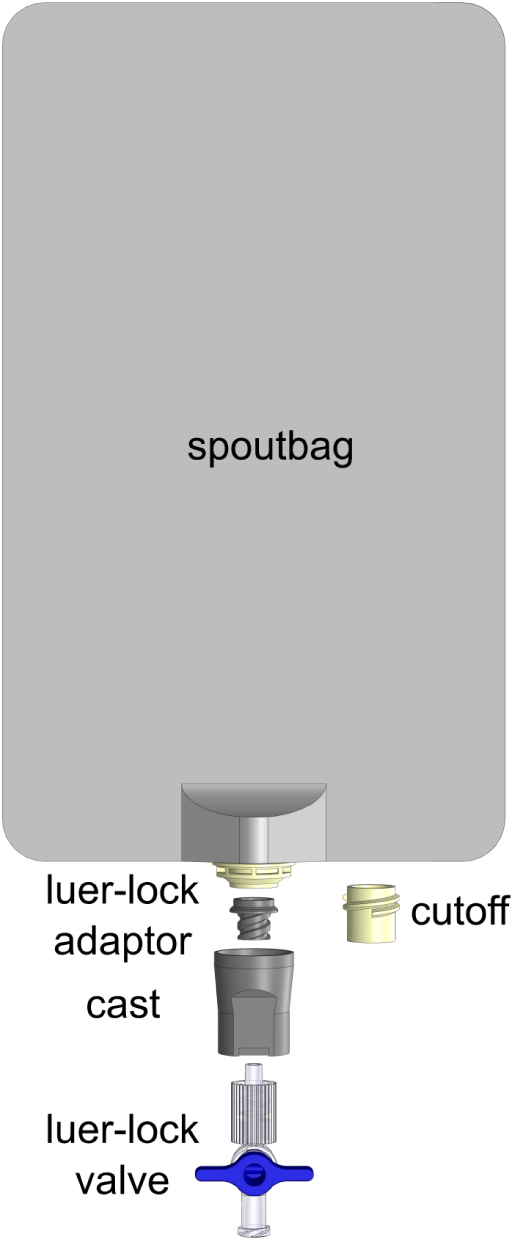
Exploded view of the gasbag assembly.

#### Synthesis of 3-nitrooxypropanol (3-NOP)

A solution of 3-bromo-1-propanol (2 g, 1.3 mL, 14.4 mmol, 1 eq) in acetonitrile (10 mL) was added to a solution of AgNO_3_ (silver nitrate) (3.67 g, 21.6 mmol, 1.5 eq) in acetonitrile (20 mL) and stirred at room temperature for 24 h. To the reaction mixture was added brine (150 mL) and stirred for 1h. The silver salts were filtered off through Celite and the filtrate was extracted with diethylether (3 × 100 mL). The organic layer was washed with brine, dried over disodium sulfate, filtered, concentrated and dried under vacuum to give the product as a yellow liquid (0.847 g, 6.99 mmol, 49%).

The analysis of 3-NOP was conducted via 1H and 13C nuclear magnetic resonance, high resolution mass spectrometry and infrared spectroscopy (IR) spectrometry and the corresponding results can be found in the supplementary data. The product was found to be of high purity with only minor acetonitrile and water residues.

#### Data evaluation

Raw data processing was done in Libre Office calculator. To identify outliers, a Grubbs test was performed. Values which had two or more consecutive outliers were then excluded from further analysis. To evaluate the variances, a one- or two-sided Student’s t-test, also known as Welch test, was performed as appropriate (64). *H_0_* was rejected if two or more timepoints in succession were significantly different, with a threshold for significance of p=0.05 for the one-tailed t-test and p=0.025 for the two-tailed t-test. Graphs of the generated data were made in GraphPadPRISM 5, figures were edited in Inkscape 1.2. For the microbiome analysis a Mann-Whitney-U-test was performed to assess whether the control was different from the stimulation (65). Protein visualisation was done with ChimeraX (56).

## Anticipated results

To facilitate basic research on methanogenesis in ruminants and to provide a tool for the development of state-of-the-art anti-methanogenic compounds, we designed an incubation setup for daily anaerobic feeding and sampling. The setup also allows for the conduction of tests on gas production and microbiome sampling. To demonstrate the possibility to cultivate anaerobic cultures for extended periods, weeks-long *in vitro* studies were performed. In this section, we demonstrate a comparison of the anti-methanogenic effects of the additives AT powder, BF, IF, 3-NOP, rapeseed oil (NC_OIL_), as well as a BF solution in oil (BF_OIL_). The investigation also encompassed an analysis of these agents’ influence on various other parameters in comparison to a negative control (NC). For each run of replicate experiments, all compounds were tested on cultures from a single inoculum prepared for each run by mixing equal parts of rumen contents from four animals. Four separate runs of experiments were conducted to produce data replicas.

### Effect of additives on total gas production

We first checked whether anti-methanogenic compounds directly affect the total gas production. The produced gas volumes were summed for those days of the experiment after stimulation where the compounds produced CH_4_ reductions and compared to the gas production of the NC on these days. The results demonstrated that 3 mg and 6 mg AT both resulted in a slight increase of 5.73% (p >0.05) and 5.19% (p >0.05) in gas production, respectively, whereas 9 mg AT resulted in a significant reduction of 17.05% (p =0.0006). 2.5 µM and 3.75 µM BF yielded significant reductions of 15.38% (p =0.002) and 9.90% (p =0.004), respectively, while 2.5 µM and 3.75 µM IF resulted in significant reductions of 12.46% (p =0.001) and 13.03% (p =0.002), respectively. The 1.25 µM BF_OIL_ treatment resulted in a reduction of 10.18% (p =0.033) and 8.64% (p =0.038) in comparison to the NC and NC_OIL_ controls, respectively. No further significant reductions in total gas production were observed.

### Effect of additives on methane production

For assessment of anti-methanogenic activity, averages of all daily produced CH_4_ volumes were calculated. The CH_4_ production, with the standard error of the mean, normalized on the corresponding negative control, is shown in Figure 5. All treatments showed a significant reduction in the CH_4_ production rate compared to the corresponding negative control. The CH₄ reduction varied depending on the compound and its respective concentration. Table 2 provides an overview of the results. Higher doses of AT, BF, and IF led to stronger and prolonged reductions. AT, BF and IF treatments at the highest concentrations achieved maximum CH₄ reductions of 96.29% on day 5 (p =5.01E-06), 98.22% on day 6 (p =2.88E-05) and 96.26% on day 5 (p =1.03E-05), respectively, all maintaining significant inhibition for 7 days (p =4.34E-05), (p =2.11E-05) and (p =0.003), respectively. No significant differences (p >0.05) were observed between BF and IF at the same concentrations. For 3-NOP, CH_4_ inhibition was moderate, with a maximum reduction of 74.63% on day 2 at 33 µM (p =8.88E-06), lasting up to 7 days (p =0.16). Lower concentrations (8.25 µM and 16.5 µM) resulted in weaker effects, with CH₄ reductions of 50.62% on day 2 (p =0.009) and 58.02% on day 2 (p =0.003), respectively. The impact of rapeseed oil alone was found to be limited, with a mere 28.95% (p =0.001) reduction in CH₄. However, BF dissolved in rapeseed oil suppressed CH_4_ production effectively. Concentration of 1.25 µM and 2.5 µM BF in oil achieved reductions of 98.51% (p =4.18E-06) and 98.02% (p =5.2E-06), respectively, and persisted for the full 7-day incubation period (p =0.0003). Table 2 also shows the day of maximum inhibition following inhibition to quantify the time delay until start of a recovery of CH₄ after stimulation.

**Figure 5:**
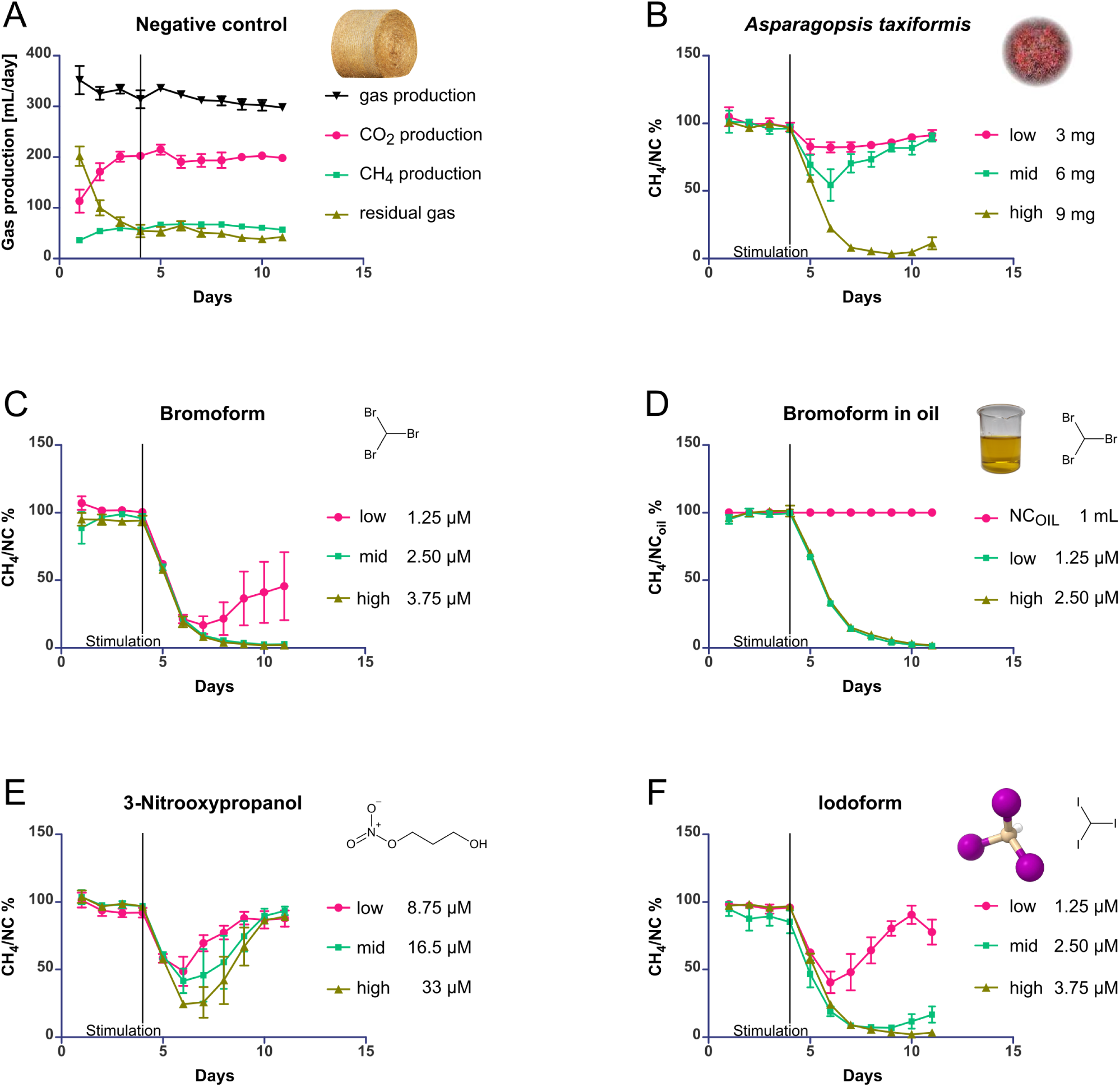
Comparison of efficacy of different anti-methanogenic additives. (A) Gas production in the negative control. Produced CH_4_ volumes for the stimulation experiments (B-F) are normalized by CH_4_ volumes produced in simultaneously conducted negative control experiments. Tested additives are Asparagopsis taxiformis (AT), bromoform (BF), rapeseed oil (NC_OIL_) and bromoform in rapeseed oil (BF_OIL_), 3-nitrooxypropanol (3-NOP), iodoform (IF); (B)-(F) The pink circles represent the lowest inhibitor concentration, the green squares the medium concentration and the brown triangles the highest inhibitor concentration. Error bars represent ± SEM; n=4 experiment repetitions.

**Table 2:** Effect of stimulation with different anti-methanogenic additives on the production of CH_4_. The reductions are normalized to the corresponding NC and given in percent of the NC. The duration of CH_4_ reduction represents the number of days following stimulation during which CH₄ production was reduced.

| Compound | Maximum $\text{CH}_4$ reduction | Duration of $\text{CH}_4$ reduction (days) | Day of Maximum Inhibition |
| --- | --- | --- | --- |
| AT |  |  |  |
| 3 mg | 17.67% | 6 | 2<sup>nd</sup> |
| 6 mg | 46.07% | 7 | 2<sup>nd</sup> |
| 9 mg | 96.29% | 7 | 5<sup>th</sup> |
| BF | 7 d |  |  |
| 1.25 $\mu\text{M}$ | 83.24% | 6 | 3<sup>rd</sup> |
| 2.5 $\mu\text{M}$ | 97.39% | 7 | 6<sup>th</sup> & 7<sup>th</sup> |
| 3.75 $\mu\text{M}$ | 98.22% | 7 | 6<sup>th</sup> |
| IF |  |  |  |
| 1.25 $\mu\text{M}$ | 59.25% | 5 | 2<sup>nd</sup> |
| 2.5 $\mu\text{M}$ | 92.85% | 7 | 4<sup>th</sup> & 5<sup>th</sup> |
| 3.75 $\mu\text{M}$ | 96.26% | 7 | 5<sup>th</sup> |
| 3-NOP |  |  |  |
| 8.25 $\mu\text{M}$ | 50.62% | 4 | 2<sup>nd</sup> |
| 16.5 $\mu\text{M}$ | 58.03% | 6 | 2<sup>nd</sup> |
| 33 $\mu\text{M}$ | 74.63% | 7 | 2<sup>nd</sup> & 3<sup>rd</sup> |
| $\text{NC}_{\text{OIL}}$ | | | |
| 1 mL | 28.96% | 7 | 4<sup>th</sup> |
| $\text{BF}_{\text{OIL}}$ | | | |
| 1.25 $\mu\text{M}$ | 98.51% | 7 | 7<sup>th</sup> |
| 2.5 $\mu\text{M}$ | 98.02% | 7 | 7<sup>th</sup> |

### Effect of additives on carbon dioxide production

Using spectroscopic data, the averages of the daily produced CO_2_ volumes were calculated. Slight, insignificant (p >0.025) increases were recorded for 9 mg AT and 3.75 µM BF with 13.46% and 7.18% respectively, as well as a significant decrease of 9.07% (p =0.009) in CO_2_ production occurred for 33 µM 3-NOP. In contrast, a significant increase in CO_2_ production was observed for 2.5 µM BF, 3.75 µM IF and 1 mL rapeseed oil, with values of 12.43% (p =0.002) 16.70% (p =0.004) and 8.8% (p =0.013), respectively.

### Effect of additives on the residual gas composition

The produced volumes of residual gas were calculated on a daily basis along with their average concentration. Given the initial pure nitrogen atmosphere and its volumetric dilution by the produced gassed, only the values beginning with the day following the stimulation were considered. The stimulations with 9 mg AT, 1.25 µM BF, 2.5 BF, 3.75 µM BF, 2.5 µM IF, 3.75 µM IF and 1.25 µM BF and 2.5 µM BF in rapeseed oil resulted in a significant increase in IR-silent molecular species, as shown in Table 3. We consider it likely that the residual gas is mainly H_2_.

**Table 3:** Effect of the stimulation with different anti-methanogenic additives on the production of residual gases. The increases are normalized to the corresponding NC and given in percent of the NC. Significance levels were calculated with two-sided t-tests.

| Compound | 9 mg AT | 1,25 μM BF | 2,5 μM BF | 3,75 μM BF | 2,5 μM IF | 3,75 μM IF | 1,25 μM BF Oil | 2,5 μM BF Oil |
| --- | --- | --- | --- | --- | --- | --- | --- | --- |
| Residual gas (likely H<sub>2</sub>) | 186.81% | 172.84% | 181.60% | 215.97% | 218.83% | 172.34% | 188.77% | 181.48% |
| Significance level | P=0.006 | P =0.024 | P =0.014 | P=0.008 | P=0.009 | P=0.009 | P=0.003 | P =0.004 |

### Repeated stimulation over 23 days of cultivation

To provide an initial assessment of reversibility of the methanogenesis inhibition, a repeated stimulation experiment was conducted. For simplicity, we extended the duration of the fourth repetition of the experiments used to accumulate statistical data for this work. The cultivation period was extended to a total duration of 23 days. After a brief regeneration phase after the first stimulation, a second stimulation was performed on day 16 of the experiment. The result was an effect similar to the first stimulation on day 4. For almost all additives, CH₄ had recovered and decreased again after the second stimulation. On average, CH₄ inhibition was 6% lower than after in the initial stimulation. The exception were BF_OIL_ treated samples, which did not recover their CH_4_ production after the initial stimulation.

### Microbiome analysis

For an evaluation of the stability of the microbiome with and without the influence of anti-methanogenic additives, the relative microbial abundance was assessed via 16S RNA metagenomics. Samples were taken prior to stimulation on day 2 and 4. Subsequently, on day 8 where the compounds showed activity, and on day 11, where CH_4_ production was typically recovering. The most prevalent taxa found in our samples were *Bacteroidetes, Euryarchaeota, Fibrobacteres, Firmicutes, Proteobacteria, Spirochaeta* and *Verrucomicrobia*. The relative abundance of the major taxa is shown in Figure 6. Taxa that were present in smaller numbers were grouped as Other, comprising *Actinobacteriota, Bdellovibrionota, Campilobacterota, Cyanobacteria, Chloroflexi, Desulfobacterota, Elusimicrobiota, Halobacterota, Patescibacteria, Planctomycetota, Synergistota,* and *WPS-2*. All unclassified counts were combined into a single category, designated as Unclassified. The complete list can be found in the supplementary data.

**Figure 6:**
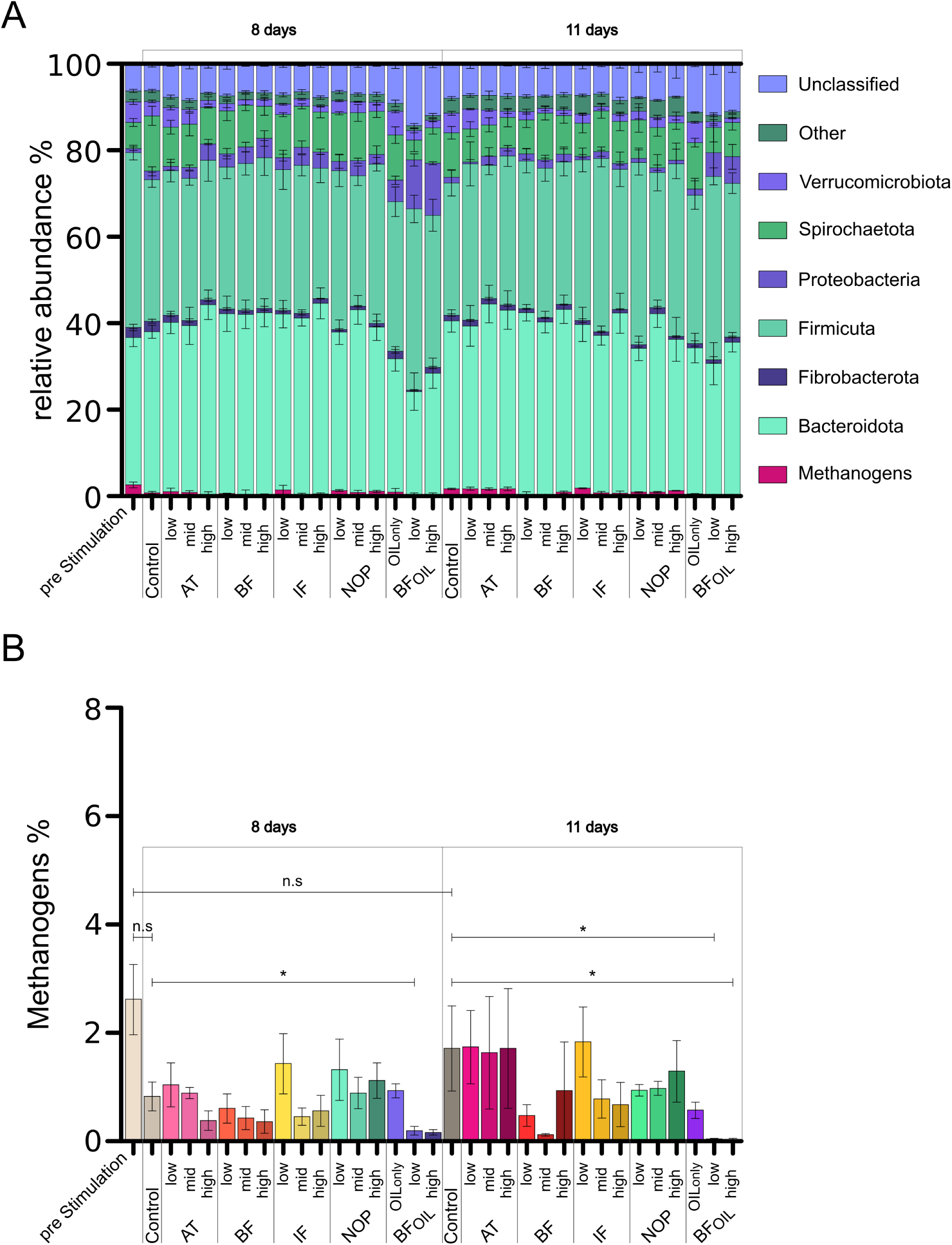
Comparison of relative abundance of microbial taxa. (A) Relative abundance of microbial taxa at the phylum level. (B) Relative abundance within the Methanogens/the Euryarchaeota phylum. BF: bromoform, IF: iodoform, 3-NOP: 3-nitrooxypropanol, BF_OIL_: bromoform in rapeseed oil. Error bars represent ± SEM; n=4 experiment repetition. Asterisk “*” indicates p <0.05.

The taxa of *Bacteroidetes* and *Firmicutes* exhibited a stable abundance, with a mean of 38% and 35%, respectively. *Verrucomicrobia*, Other and Unclassified had mean abundances of 4%, 3%, and 6% respectively. *Fibrobacteres* were not present in run 4 and therefore treated as outliers. *Fibrobacteres* showed an expression of 1%-3%. The population of *Proteobacteria* exhibited an increase from day 4 to day 8, followed by a return to baseline levels after 11 days. This population ranged from <1% to 5% with the exception of the BFOIL stimulation, where the abundance increased to 6%. The population of *Spirochaeta* began the experiment at 3% and increased its abundance to 9% at the end of the experiment. Following an initial adaptation phase, the methanogens exhibited an abundance comparable to that observed in the pre-stimulation sample, with the exception of the data for BFOIL stimulation. The classes of *Methanomassiliicoccales, Methanobrevibacter, Methanosphera* and *Thermoplasmata* were the reported order or genus of the Methanogens or *Euryarchaeota* present in the samples. Their abundance was reduced with the progression of the incubation from 3% to <1% at day 8 and back to up to 2% at day 11. The greatest reduction in Methanogens was observed in the BFOIL stimulation. The taxon of *Bacteroidetes* did not differ significantly (p >0.025) from the negative control in all samples. *Fibrobacteres* exhibited significant (p =0.025) differences in the 8.75 µM, 16.5 µM, and 33 µM 3-NOP stimulations, and the NC_OIL_ after 8 days, resulting in mean reductions of 76%, 66%, 67%, and 88%, respectively. *Firmicutes* were significantly (p =0.025) increased in the 1.25 µM BF_OIL_ stimulation with an average of 39%. Methanogens showed a significant (p =0.025) reduction in relative abundace in the 2.5 µM BF_OIL_ set after 8 days, and in the 1.25 µM and 2.5 µM BF_OIL_ groups after 11 days, with 80%, 98%, and 98% respectively. The *Proteobacteria* showed no significant (p >0.025) differences compared to the negative control. *Spirochaeta* were significantly reduced (p =0.025) for the 1.25 µM BF_OIL_ stimulation of four days with 64%. *Verrucomicrobia* exhibited considerable decreases for 9 mg AT, for 1.25 µM and 2.5 µM BF, and 3.75 µM IF after 8 days, with reductions of 52%, 50%, 61% and 57%, respectively. After 11 days, significant differences in abbundances of *Verrucomocrobia* were observed (p =0.025) with 9 mg AT, 1.25 µM, 2.5 µM and 3.75 µM BF and IF, 8.75 µM, 16.5 µM and 33 µM 3-NOP and 1.25 µM and 2.5 µM BF_OIL_, with reductions of 68%, 57%, 78%, 72%, 55%, 79%, 64%, 56%, 49%, 66%, 71%, and 79%, respectively. The abundances of other taxa were significantly different (p =0.025) for the 1.25 µM and 2.5 µM BF_OIL_ groups after 8 and 11 days with a reductions of 53% and 57%, and 59% and 57%, respectively. Unclassified counts were significantly (p =0.025) elevated for 1.25 µM BF_OIL_ after 8 and 11 days, with 129% and 50% respectively. Over the course of the experiment, the negative controls showed a increase solely in *Spirochaeta* between days 4 and 8, with an increase of 111% (p =0.025).

## Discussion

Although methanogenesis in the digestive system of ruminants is an active research area due to its potential impact on global warming (17,66,67), the establishment of a robust, reliable and affordable *in vitro* rumen cultivation system remains challenging in laboratories where expensive equipment is unavailable. Such setups are, however, required to investigate a multitude of rumen fermentation parameters over the course of time. With the development of our setup, we sought to combine the advantages of the two best-established systems, namely the throughput and ease-of-use of batch systems and the quality and stability of rumen simulation setups. The setup was subjected to thorough tests and we demonstrated its use by comparing a selection of anti-methanogenic additives.

The results demonstrate the stability of the microbiome and the consistency of the overall gas production rates and the production rates of individual gas species. Experiments were routinely conducted for 11 days and in one case even for 23 days. Thus, the easily replicable, self-made setup is suitable for rumen cultivation experiments over days and even weeks. The stability of the predominant phyla *Bacteriodota* and *Firmicutes*, along with the comparable abundance of *Fibrobacteres*, suggests that this setup exhibits similar properties to state-of-the-art cultivation systems, although the vast variance of the inocula and the different PCR-primers make a comparison with published data difficult (68,69). The average gas production of 165.44 mL/day per gram of daily fed dry matter (gDM/day) in this setup is comparable to the daily gas production of other batch cultures and RUSITEC setups (28,40,45). However, there are minor discrepancies with some published figures. In detail, we measured a 9% higher daily gas production than reported for the batch cultivation system of Machado *et al.* (45) Also, a 26% higher gas production compared to the initial RUSITEC setup from Czerkawski and Breckenridge (40) was found, which is not unexpected because the liquid turnover in our setup is considerably smaller than in RUSITEC setups. Differences in gas production between separate studies may also be due to different inocula, different substrates, and differences in nutrient addition procedures, where, for example, feed may accumulate in the cultures over time if the degradation rate is slow. Also, our culture bottles contained ceramic cylinders that can allow bacteria to attach and help to retain the solid fraction. The potential direct effects of the cylinders, as well as any microbes attached to them, remain to be evaluated in the future.

For this study, we drew our inoculum from randomly selected rumen contents of animals of different sex, breed, diet, health status and potential previous antibiotic treatments. Consequently, the variability of the microbiome is pronounced and represents a sample of the cattle population raised in Bavaria, Germany. All additives were tested on all rumen cultures and the negative control cultures came from the same animals as the cultures used for stimulation experiments. However the initial rumen culture compositions necessarily varied between repetitions of the experiments. As would be expected, a large variability was found in the reported amount of the initial fraction of *Euryarchaeota* (68,70,71). After an adaptation period after stimulation with most anti-methanogenic additives, the abundance of methanogens did not differ significantly (p >0.05) from that observed in the negative controls. The notable exception is stimulation with bromofom in oil (BF_OIL_), where a substantial and apparently irreversible reduction of methanogen abundance was observed. For this additive, results of the CH_4_ production and the microbiome analysis are concordant. It is important to note that the sampling process did not consider the attachment of certain bacteria to particles or the ceramic cylinders. It was observed that the overall abundance of methanogens varied also in the negative controls, in particular during the initial culture start-up phase, although the CH_4_ production remained markedly stable. Improved homogenisation of the samples may help to reduce the large variations observed. For a more detailed analysis of the microbiom’s response, the analysis can be refined with complementary methods, for example, for evaluation of protozoal counts. Primers for the replication of the 16sRNA fragments can also be optimized for better resolution.

The demonstration experiments with our setup provide a valuable direct comparison of the potency of different CH_4_ mitigation strategies, including the supplementation of 3-NOP, AT, BF, IF and BF in an oil solution. The reductions of CH_4_ production after addition of AT and BF are consistent with values reported in the literature (68). Moreover, we found that 3-NOP consistently exhibited a weaker effect on methanogenesis than BF or IF. However, our results indicate a slightly better anti-methanogenic effect of 3-NOP than expected from literature values. Previously, a 76% reduction of CH_4_ production has been reported for a supplementation of approximately 41.3 µM of 3-NOP (47). In our experiments, 33 µM of 3-NOP were administered once to achieve a maximum reduction of 74.63% of CH_4_ production. Stimulation with 3-NOP yielded the most transient effect, with CH_4_ production returning to its previous levels after a period of only five days.

In conclusion, the setup and protocols developed, together with the demonstration experiments performed, provide a solid methodological basis for future *in vitro* studies of methanogenesis. Such experimental approaches greatly facilitate the first step towards the development of anti-methanogenic additives. Moreover, the approaches may contribute to a better understanding of some of the controversial results from *in vivo* experiments in cattle (27,29,31,72,73), which suggest a need for further research in this area.

## Acknowledgements

We thank Volta Greentech for the kind gift of the red algae, *Asparagopsis taxiformis*. We thank the staff members of Attenberger Fleisch GmbH & Co. KG Slaughterhouse in Munich for provision of samples and an introduction to probing the rumen. We are grateful to Dr. Markus Müller from the Institute for Chemical Epigenetics at LMU Munich for support of our compound synthesis.

## Supplements

**Figure 7:**
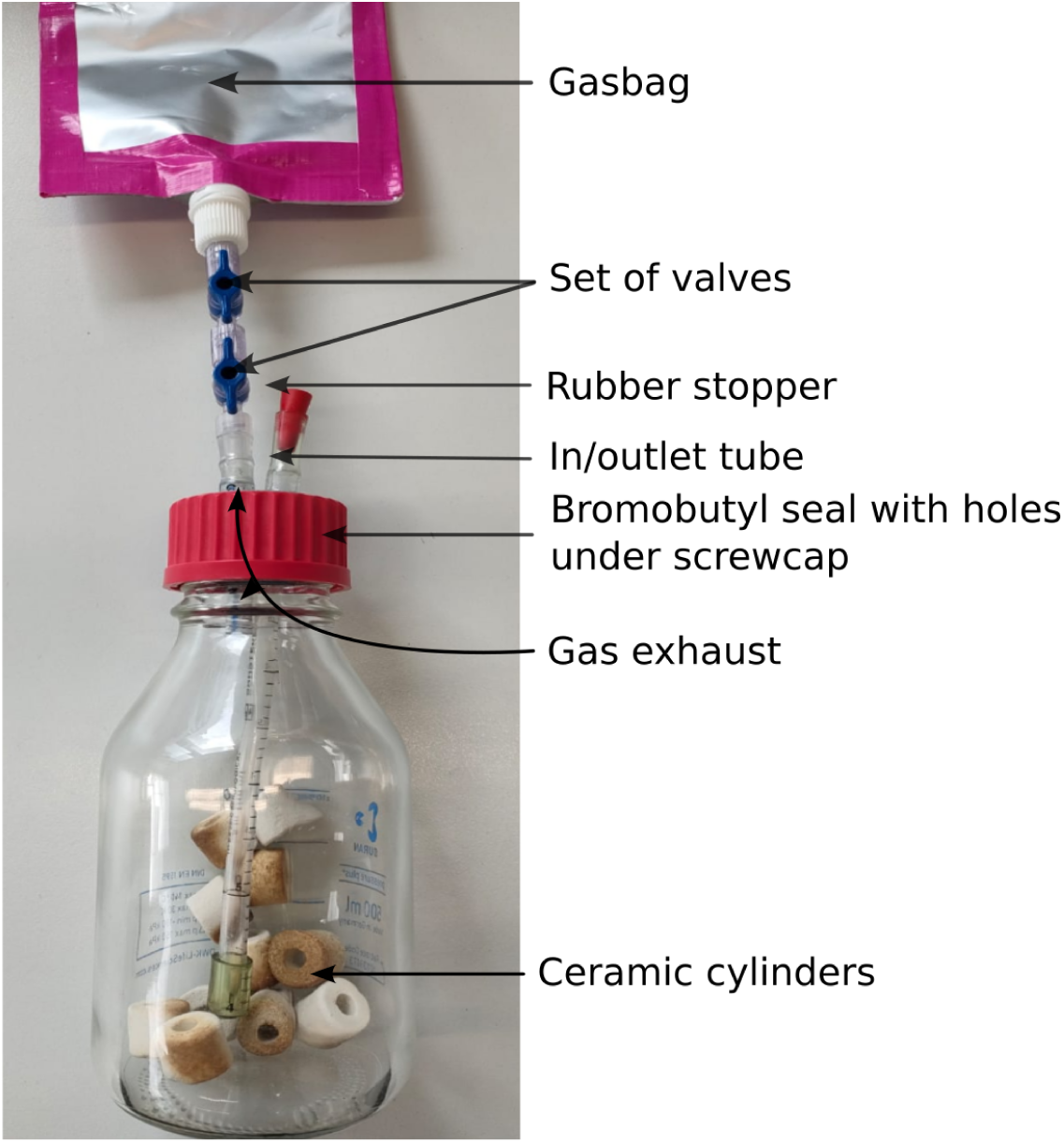
Prototype of the easily feedable rumen incubation flask. From top to bottom consisting of a reinforced 500 mL spout bag, Luer-lock connectors and valves, two short PVC tubes mounted on serological pipettes, ceramic cylinders and the 500 mL glass bottle with bromobutyl seal and screw cap.

**Table 4:**
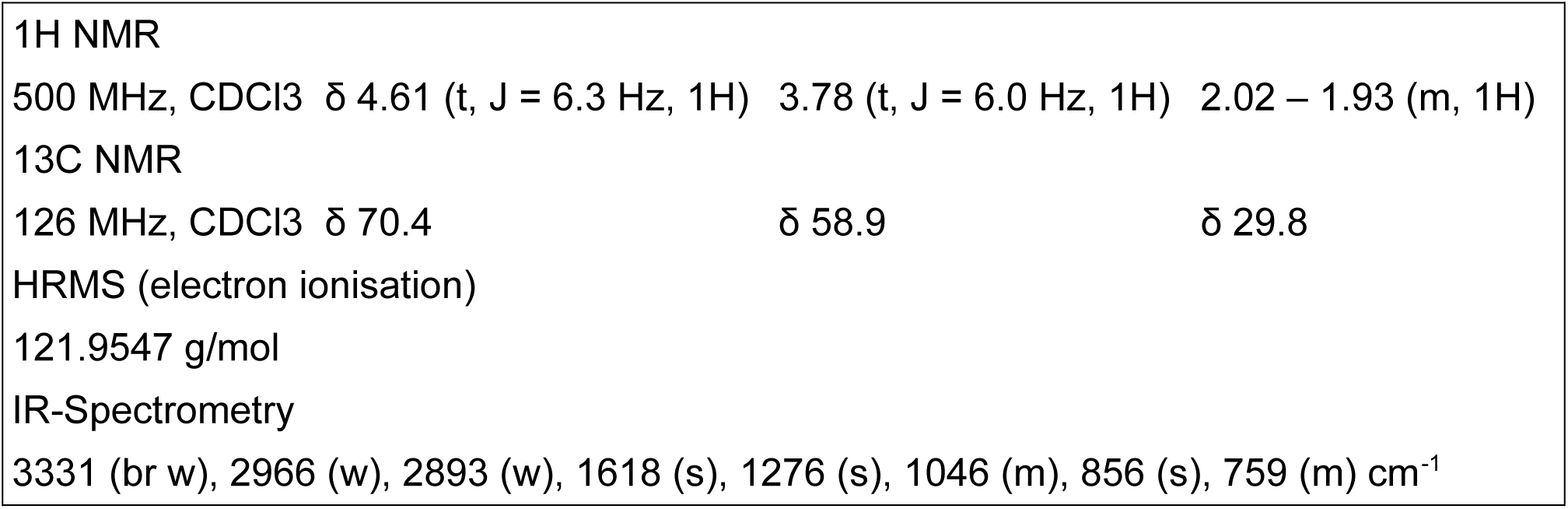
Analysis of 3-nitrooxypropanol by 1H NMR, 13C NMR, HRMS and IR spectrometry Evaluation of the synthesised 3-NOP by NMR, HRMS and IR spectrometry shows low acetonitril impurities at δ 2.09 and of water at δ 1.59.

**Table 5.**
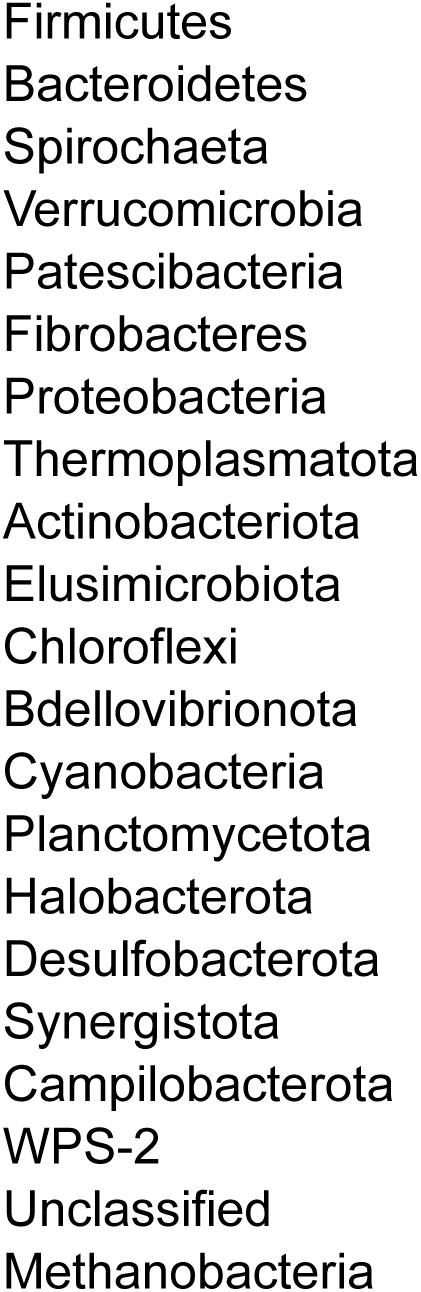
List of reported taxa with 16S RNA sequencing In the above figure, in contrast to Fig. 4 in the paper, you can see the initial acclimation period. Although the initial abundance of methanogenic *Archaea* is higher, the methane production is not. Another noteworthy point is, that the high variance is due to the blinded sampling, which is in contrast to the well defined inoculi of other studies.

**Figure 8:**
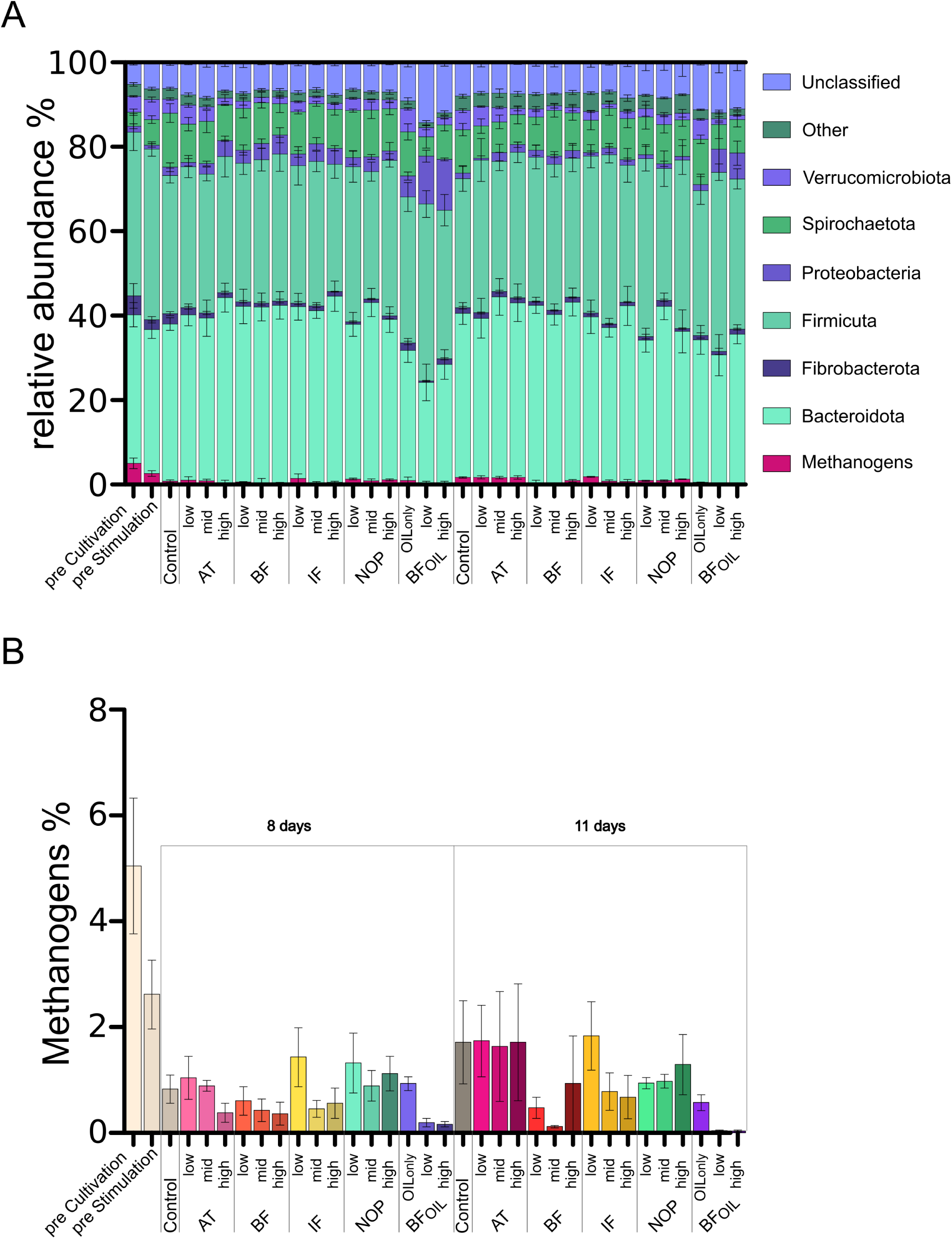
Relative abundance of microbial taxa as determined by 16S rRNA amplicon sequencing over a course of 11 days. Error bars represent ± SEM (n=4) (A) Relative abundance of microbial taxa at the phylum level. (B) Relative abundance within the Methanogens. the Euryarchaeota phylum. BF: bromoform, IF: iodoform, 3-NOP: 3-nitrooxypropanol, BFOIL: bromoform in Oil.

## References

1. Cedervall PE, Dey M, Li X, Sarangi R, Hedman B, Ragsdale SW, et al. Structural Analysis of a Ni-Methyl Species in Methyl-Coenzyme M Reductase from Methanothermobacter marburgensis. J Am Chem Soc. 2011 Apr 20;133(15):5626–8.

2. United Nations. Adoption of the Paris Agreement [Internet]. 2015 [cited 2024 Jun 14]. Available from: https://unfccc.int/sites/default/files/l09r01.pdf

3. Climate Watch [Internet]. 2019 [cited 2023 Apr 19]. Available from: https://www.climatewatchdata.org/sectors/agriculture?emissionType=203&filter=

4. Dean J. Methane, climate change, and our uncertain future. Eos. 2018 May 11;99.

5. Stevenson DS, Zhao A, Naik V, O’Connor FM, Tilmes S, Zeng G, et al. Trends in global tropospheric hydroxyl radical and methane lifetime since 1850 from AerChemMIP. Atmospheric Chem Phys. 2020 Nov 5;20(21):12905–20.

6. Nisbet-Jones PBR, Fernandez JM, Fisher RE, France JL, Lowry D, Waltham DA, et al. Is the destruction or removal of atmospheric methane a worthwhile option? Philos Trans R Soc Math Phys Eng Sci. 2021 Dec 6;380(2215):20210108.

7. Ripple WJ, Smith P, Haberl H, Montzka SA, McAlpine C, Boucher DH. Ruminants, climate change and climate policy. Nat Clim Change. 2014 Jan;4(1):2–5.

8. Gerber PJ, FAO, editors. Tackling climate change through livestock: a global assessment of emissions and mitigation opportunities. Rome: Food and Agriculture Organization of the United Nations; 2013. 115 p.

9. Pachauri RK, Reisinger A, IPCC, editors. Climate change 2007: Synthesis report: contribution of Working Groups I, II and III to the Fourth Assessment Report of the Intergovernmental Panel on Climate Change. Geneva: IPCC; 2008. 103 p.

10. Akin DE, Benner R. Degradation of polysaccharides and lignin by ruminal bacteria and fungi. Appl Environ Microbiol. 1988 May;54(5):1117–25.

11. Wolin MJ. Fermentation in the rumen and human large intestine. Science. 1981 Sep 25;213(4515):1463–8.

12. Wang M, Janssen PH, Sun XZ, Muetzel S, Tavendale M, Tan ZL, et al. A mathematical model to describe in vitro kinetics of H2 gas accumulation. Anim Feed Sci Technol. 2013 Aug 9;184(1):1–16.

13. Ungerfeld EM. Shifts in metabolic hydrogen sinks in the methanogenesis-inhibited ruminal fermentation: a meta-analysis. Front Microbiol. 2015;6.

14. Zheng Y, Kahnt J, Kwon IH, Mackie RI, Thauer RK. Hydrogen formation and its regulation in Ruminococcus albus: involvement of an electron-bifurcating [FeFe]-hydrogenase, of a non-electron-bifurcating [FeFe]-hydrogenase, and of a putative hydrogen-sensing [FeFe]-hydrogenase. J Bacteriol. 2014 Nov;196(22):3840–52.

15. Johnson KA, Johnson DE. Methane emissions from cattle. J Anim Sci. 1995 Aug 1;73(8):2483–92.

16. Dodds KG, Auvray B, Newman SAN, McEwan JC. Genomic breed prediction in New Zealand sheep. BMC Genet. 2014 Sep 16;15(1):92.

17. Almeida AK, Hegarty RS, Cowie A. Meta-analysis quantifying the potential of dietary additives and rumen modifiers for methane mitigation in ruminant production systems. Anim Nutr. 2021 Dec 1;7(4):1219–30.

18. Bauchop T. Inhibition of Rumen Methanogenesis by Methane Analogues. J Bacteriol. 1967 Jul;94(1):171–5.

19. Lanigan GW. Metabolism of pyrrolizidine alkaloids in the ovine rumen. II. Some factors affecting rate of alkaloid breakdown by rumen fluid in vitro. Aust J Agric Res. 1970;21(4):633–40.

20. Lanigan GW, Smith LW. Metabolism of pyrrolizidine alkaloids in the ovine rumen. I. Formation of 7-α-hydroxy-1-α-methyl-8-α pyrrolizidine from heliotrine and lasiocarpine. Aust J Agric Res. 1970;21(3):493–500.

21. Van Nevel CJ, Demeyer DI. Effect of Methane Inhibitors on the Metabolism of Rumen Microbes in Vitro. Arch Für Tierernaehrung. 1981 Feb 1;31(2):141–51.

22. Lanigan GW. Metabolism of pyrrolizidine alkaloids in the ovine rumen. III. The competitive relationship between Heliotrine metabolism and methanogenesis in the rumen fluid in vitro. Aust J Agric Res. 1971;22(1):123–30.

23. Lanigan GW. Metabolism of pyrrolizidine alkaloids in the ovine rumen. IV. Effects of chloral hydrate and halogenated methanes on rumen methanogenesis and alkaloid metabolism in fistulated sheep. Aust J Agric Res. 1972;23(6):1085–91.

24. Trei JE, Parish RC, Singh YK, Scott GC. Effect of Methane Inhibitors on Rumen Metabolism and Feedlot Performance of Sheep. J Dairy Sci. 1971 Apr 1;54(4):536–40.

25. Czerkawski JW, Breckenridge G. New inhibitors of methane production by rumen micro-organisms. Development and testing of inhibitors in vitro. Br J Nutr. 1975 Nov;34(3):429–46.

26. Johnson DE. Effects of a Hemiacetal of Chloral and Starch on Methane Production and Energy Balance of Sheep Fed a Pelleted Diet. J Anim Sci. 1972 Nov 1;35(5):1064–8.

27. Thorsteinsson M, Lund P, Weisbjerg MR, Noel SJ, Schönherz AA, Hellwing ALF, et al. Enteric methane emission of dairy cows supplemented with iodoform in a dose– response study. Sci Rep. 2023 Aug 7;13:12797.

28. Machado L, Magnusson M, Paul NA, Nys R de, Tomkins N. Effects of Marine and Freshwater Macroalgae on In Vitro Total Gas and Methane Production. PLOS ONE. 2014 Jan 22;9(1):e85289.

29. Roque BM, Venegas M, Kinley RD, Nys R de, Duarte TL, Yang X, et al. Red seaweed (*Asparagopsis taxiformis*) supplementation reduces enteric methane by over 80 percent in beef steers. PLOS ONE. 2021 Mar 17;16(3):e0247820.

30. Glasson CRK, Kinley RD, de Nys R, King N, Adams SL, Packer MA, et al. Benefits and risks of including the bromoform containing seaweed Asparagopsis in feed for the reduction of methane production from ruminants. 2022 May [cited 2022 Jun 10]; Available from: https://hdl.handle.net/10182/14753

31. Romero P, Belanche A, Jiménez E, Hueso R, Ramos-Morales E, Salwen JK, et al. Rumen microbial degradation of bromoform from red seaweed (*Asparagopsis taxiformis*) and the impact on rumen fermentation and methanogenic archaea. J Anim Sci Biotechnol. 2023 Nov 1;14(1):133.

32. Jaun B, Thauer R. Methyl-Coenzyme M Reductase and its Nickel Corphin Coenzyme F430 in Methanogenic Archaea. 2007 Mar 1;323–56.

33. Duin EC, Wagner T, Shima S, Prakash D, Cronin B, Yáñez-Ruiz DR, et al. Mode of action uncovered for the specific reduction of methane emissions from ruminants by the small molecule 3-nitrooxypropanol. Proc Natl Acad Sci. 2016 May 31;113(22):6172–7.

34. Jin W, Meng Z, Wang J, Cheng Y, Zhu W. Effect of Nitrooxy Compounds with Different Molecular Structures on the Rumen Methanogenesis, Metabolic Profile, and Methanogenic Community. Curr Microbiol. 2017 Aug;74(8):891–8.

35. Muetzel S, Hunt C, Tavendale MH. A fully automated incubation system for the measurement of gas production and gas composition. Anim Feed Sci Technol. 2014 Oct 1;196:1–11.

36. Muetzel S, Ronimus RS, Lunn K, Kindermann M, Duval S, Tavendale M. A small scale rumen incubation system to screen chemical libraries for potential methane inhibitors. Anim Feed Sci Technol. 2018 Oct 1;244:88–92.

37. Durmic Z, Duin EC, Bannink A, Belanche A, Carbone V, Carro MD, et al. *Feed additives for methane mitigation:* Recommendations for identification and selection of bioactive compounds to develop antimethanogenic feed additives. J Dairy Sci. 2025 Jan 1;108(1):302–21.

38. Soto EC, Molina-Alcaide E, Khelil H, Yáñez-Ruiz DR. Ruminal microbiota developing in different in vitro simulation systems inoculated with goats’ rumen liquor. Anim Feed Sci Technol. 2013 Sep 23;185(1):9–18.

39. Mansfield HR, Endres MI, Stern MD. Comparison of microbial fermentation in the rumen of dairy cows and dual flow continuous culture. Anim Feed Sci Technol. 1995 Sep 1;55(1):47–66.

40. Czerkawski JW, Breckenridge G. Design and development of a long-term rumen simulation technique (Rusitec). Br J Nutr. 1977 Nov;38(3):371–84.

41. Hoover WH, Crooker BA, Sniffen CJ. Effects of Differential Solid-Liquid Removal Rates on Protozoa Numbers in Continous Cultures of Rumen Contents. J Anim Sci. 1976 Aug 1;43(2):528–34.

42. Abe M, Kumeno F. In Vitro Simulation of Rumen Fermentation: Apparatus and Effects of Dilution Rate and Continuous Dialysis on Fermentation and Protozoal Population. J Anim Sci. 1973 May 1;36(5):941–8.

43. Wood JM, Kennedy FScott, Wolfe RS. Reaction of multihalogenated hydrocarbons with free and bound reduced vitamin B12. Biochemistry. 1968 May 1;7(5):1707–13.

44. Paul N, de Nys R, Steinberg P. Chemical defence against bacteria in the red alga Asparagopsis armata: Linking structure with function. Mar Ecol-Prog Ser. 2006 Jan 11;306:87–101.

45. Machado L, Magnusson M, Paul N, Kinley R, de Nys R, Tomkins N. Identification of bioactives from the red seaweed Asparagopsis taxiformis that promote antimethanogenic activity in vitro. J Appl Phycol. 2016 Oct 1;28.

46. Romero-Perez A, Okine EK, McGinn SM, Guan LL, Oba M, Duval SM, et al. The potential of 3-nitrooxypropanol to lower enteric methane emissions from beef cattle. J Anim Sci. 2014 Oct;92(10):4682–93.

47. Romero Pérez A, Okine E, Guan L, Duval S, Kindermann M, Beauchemin K. Effects of 3-nitrooxypropanol on methane production using the rumen simulation technique (Rusitec). Anim Feed Sci Technol. 2015 Sep 1;209:98–109.

48. Martínez-Fernández G, Abecia L, Arco A, Cantalapiedra-Hijar G, Martín-García AI, Molina-Alcaide E, et al. Effects of ethyl-3-nitrooxy propionate and 3-nitrooxypropanol on ruminal fermentation, microbial abundance, and methane emissions in sheep. J Dairy Sci. 2014 Jun 1;97(6):3790–9.

49. Reynolds CK, Humphries DJ, Kirton P, Kindermann M, Duval S, Steinberg W. Effects of 3-nitrooxypropanol on methane emission, digestion, and energy and nitrogen balance of lactating dairy cows. J Dairy Sci. 2014 Jun 1;97(6):3777–89.

50. Kinley RD, Tan S, Turnbull J, Askew S, Harris J, Roque BM. Exploration of Methane Mitigation Efficacy Using Asparagopsis-Derived Bioactives Stabilized in Edible Oil Compared to Freeze-Dried Asparagopsis in Vitro. Am J Plant Sci. 2022 Jul 11;13(7):1023–41.

51. Alvarez-Hess PS, Jacobs JL, Kinley RD, Roque BM, Neachtain ASO, Chandra S, et al. Twice daily feeding of canola oil steeped with *Asparagopsis armata* reduced methane emissions of lactating dairy cows. Anim Feed Sci Technol. 2023 Mar 1;297:115579.

52. Machado L, Magnusson ME, Tomkins NW, KINLEY RD, Nys PCD, PAUL NA. Method for reducing total gas production and/or methane production in a ruminant animal [Internet]. EP3102219B1, 2020 [cited 2025 Feb 28]. Available from: https://patents.google.com/patent/EP3102219B1/en?q=(asparagopsis+taxiformis)&oq=asparagopsis+taxiformis

53. Duval S, Kindermann M. Use of nitrooxy organic molecules in feed for reducing methane emission in ruminants [Internet]. EP2654455B1, 2018 [cited 2023 Apr 20]. Available from: https://patents.google.com/patent/EP2654455B1/en?q=(3-nitrooxypropanol+EU)&oq=3-nitrooxypropanol+EU

54. De Nys R, Magnusson M. Novel Composition. WO2020113279A1, 2020.

55. Callahan BJ, McMurdie PJ, Rosen MJ, Han AW, Johnson AJA, Holmes SP. DADA2: High-resolution sample inference from Illumina amplicon data. Nat Methods. 2016 Jul;13(7):581–3.

56. Pettersen EF, Goddard TD, Huang CC, Meng EC, Couch GS, Croll TI, et al. UCSF ChimeraX: Structure visualization for researchers, educators, and developers. Protein Sci Publ Protein Soc. 2021 Jan;30(1):70–82.

57. DSMZ. 330: RUMEN BACTERIA MEDIUM [Internet]. 2022 [cited 2025 Feb 27]. Available from: https://www.dsmz.de

58. McDougall EI. Studies on ruminant saliva. 1. The composition and output of sheep’s saliva. Biochem J. 1948;43(1):99–109.

59. Yu Z, Morrison M. Improved extraction of PCR-quality community DNA from digesta and fecal samples. BioTechniques. 2004 May;36(5):808–12.

60. 16S Illumina Amplicon Protocol : earthmicrobiome [Internet]. 2022 [cited 2022 May 13]. Available from: https://earthmicrobiome.org/protocols-and-standards/16s/

61. Parada AE, Needham DM, Fuhrman JA. Every base matters: assessing small subunit rRNA primers for marine microbiomes with mock communities, time series and global field samples. Environ Microbiol. 2016;18(5):1403–14.

62. Apprill A, McNally S, Parsons R, Weber L. Minor revision to V4 region SSU rRNA 806R gene primer greatly increases detection of SAR11 bacterioplankton. Aquat Microb Ecol. 2015 Jun 4;75(2):129–37.

63. Lee M. A full example workflow for amplicon data [Internet]. 2023 [cited 2023 Apr 26]. Available from: https://astrobiomike.github.io/amplicon/dada2_workflow_ex#dereplication

64. Welch BL. The Significance of the Difference between two Means when the Population Variances are unequal. Biometrika [Internet]. 1938 Feb 1 [cited 2023 May 2]; Available from: https://www.scinapse.io/papers/1973158119

65. Wilcoxon F. Individual Comparisons by Ranking Methods. Biom Bull. 1945;1(6):80–3.

66. Patra AK. Enteric methane mitigation technologies for ruminant livestock: a synthesis of current research and future directions. Environ Monit Assess. 2012 Apr 1;184(4):1929–52.

67. Broucek J. Options to methane production abatement in ruminants: A review. J Anim Plant Sci. 2018 Apr 1;28:348–64.

68. Machado L, Tomkins N, Magnusson M, Midgley DJ, de Nys R, Rosewarne CP. In Vitro Response of Rumen Microbiota to the Antimethanogenic Red Macroalga Asparagopsis taxiformis. Microb Ecol. 2018 Apr 1;75(3):811–8.

69. Wetzels SU, Eger M, Burmester M, Kreienbrock L, Abdulmawjood A, Pinior B, et al. The application of rumen simulation technique (RUSITEC) for studying dynamics of the bacterial community and metabolome in rumen fluid and the effects of a challenge with Clostridium perfringens. PLOS ONE. 2018 Feb 7;13(2):e0192256.

70. Ziemer CJ, Sharp R, Stern MD, Cotta MA, Whitehead TR, Stahl DA. Comparison of microbial populations in model and natural rumens using 16S ribosomal RNA-targeted probes. Environ Microbiol. 2000;2(6):632–43.

71. Mateos I, Ranilla MJ, Saro C, Carro MD. Shifts in microbial populations in Rusitec fermenters as affected by the type of diet and impact of the method for estimating microbial growth (15N v. microbial DNA). Animal. 2017 Jan 1;11(11):1939–48.

72. Kebreab E, Bannink A, Pressman EM, Walker N, Karagiannis A, Gastelen S van, et al. A meta-analysis of effects of 3-nitrooxypropanol on methane production, yield, and intensity in dairy cattle. J Dairy Sci. 2023 Feb 1;106(2):927–36.

73. Patra AK, Puchala R. Methane mitigation in ruminants with structural analogues and other chemical compounds targeting archaeal methanogenesis pathways. Biotechnol Adv. 2023 Dec 1;69:108268.

